# A low-matrix organoid-T cell co-culture platform for functional evaluation of T cell engagers in patient-derived colorectal cancer organoids

**DOI:** 10.64898/2026.09.21.751493

**Authors:** Claudia M. A. Pinna, Kamila Rakhimova, Jing Zhang, Laura Bills, Louise Tee, Charlotte Blanc, David G. Millar, Elsa Rottoli, Oisín Huhn, Miguel Gaspar, Neeraj Lal, Romain Lara, Kaitlin A. Marley, Saso Cemerski, Simon J. Dovedi, Andrew D. Beggs

**Author notes:** These authors contributed equally (Kamila Rakhimova, Jing Zhang). Senior author (Claudia M. A. Pinna).

## Abstract

**Background:** Colorectal cancer (CRC) remains a leading cause of cancer-related mortality, and emerging T cell engager (TCE) immunotherapies require predictive preclinical models that capture patient-specific tumour biology and tumour-immune interactions. Although patient-derived organoid (PDO)-immune co-culture systems show promise for evaluating immunotherapy responses, many rely on matrix-embedded cultures that limit scalability and reproducibility. We therefore developed and validated a low-matrix organoid-T cell co-culture platform for functional assessment of TCE activity in patient-derived CRC models.

**Methods:** Patient-derived CRC organoids representing diverse molecular and genetic backgrounds were co-cultured with activated allogeneic CD3+ T cells in a suspension low-matrix format. Matrix concentration, medium composition, T cell activation status, and assay duration were optimized. Tumour killing, T cells activation, cytokine secretion, apoptosis, and motility were assessed using flow cytometry, live-cell imaging, cytokine profiling, and immunofluorescence. The platform was evaluated using EGFR- and HER2-targeting bispecific T cell engagers across 21 CRC organoid models and multiple healthy donor-derived T cell populations.

**Results:** Optimization identified 1% matrix and a 1:1 organoid: T cell medium that preserved organoid integrity while maintaining T cell viability, activation, and motility. The final workflow enabled reproducible co-culture of 5-day mature organoids with 7-day activated T cells for 72 hours. EGFR-targeting TCEs induced tumour killing, T cell activation, and IFN-γ secretion across donor-organoid combinations. Screening of 21 CRC organoid models revealed substantial inter-patient heterogeneity, with 12 models maintaining ≥50% baseline viability and supporting functional TCE evaluation. EGFR- and HER2-targeting TCEs produced potent dose-dependent cytotoxicity, with IC50 values ranging from 0.0188 to 54.77 nM across responsive models and up to 75%-85% tumour killing in the most sensitive organoids. These responses were accompanied by increased effector cytokine secretion. Real-time imaging confirmed dynamic T cell engagement and apoptosis-driven organoid destruction following TCE treatment.

**Conclusions:** We establish a robust, scalable, human-relevant low-matrix organoid-T cell co-culture platform for functional screening of T cell engagers in patient-derived CRC models. By enabling integrated assessment of tumour killing, immune activation, cytokine responses, and tumour-immune interactions while preserving inter-patient heterogeneity, this system provides a translational framework for immunotherapy development. The platform may support candidate prioritisation, biomarker discovery, patient stratification, and preclinical evaluation of immune-engaging therapeutics.

## INTRODUCTION

Colorectal cancer (CRC) is the third leading cause of cancer-related mortality worldwide(1). Despite advances in surgery, chemotherapy, and radiotherapy, treatment responses remain heterogeneous and outcomes are often limited by toxicity and therapeutic resistance(2). Increasing evidence demonstrates that the tumour microenvironment (TME) is a major determinant of disease progression and therapeutic response, highlighting immunotherapy as an attractive treatment strategy for CRC(3,4).

Immunotherapy has transformed cancer treatment, and T cell-engaging bispecific antibodies are emerging as a promising strategy for redirecting cytotoxic T lymphocytes towards tumour cells(5,6). However, predictive preclinical models remain necessary to evaluate efficacy, identify responsive patient populations, and understand mechanisms of resistance(7).

There is therefore a critical need for predictive preclinical models that capture patient heterogeneity, treatment-response variability, and key features of the tumour microenvironment to bridge the translational gap between drug discovery and clinical trials.

Conventional cell-line and animal models remain important for drug development but fail to faithfully recapitulate human tumour biology, heterogeneity, and immune interactions, contributing to limited predictive performance during clinical translation(8–10).

Recent regulatory initiatives, including the FDA Modernization Act 3.0 and the Roadmap to Reducing Animal Testing in Preclinical Safety Studies, have further increased interest in human-relevant preclinical platforms, including patient-derived organoids (PDOs) and other translational *in vitro* systems(11–14).

PDOs are three-dimensional in vitro cultures derived from adult stem cells (ASCs) or pluripotent stem cells and maintained in organ-specific growth factor cocktails and extracellular matrices(15–17). PDOs recapitulate tissue architecture, preserve tumour genomic landscapes, reflect inter-patient heterogeneity, and can predict patient responses to chemotherapy and radiotherapy with high sensitivity and specificity(18–20). These features make PDOs well suited for investigating drug resistance mechanisms and evaluating therapeutic efficacy in CRC and other cancers(21–23). Importantly, PDOs retain patient-specific tumour antigens, positioning them as powerful tools for immuno-oncology research compared with murine models, which lack fully humanised immune systems and fail to capture the complexity of human tumour antigen diversity and biology(24,25).

Beyond therapeutic screening, PDO-based platforms may facilitate identification of biomarkers associated with immune-engager response and resistance, supporting patient-stratification strategies. However, PDOs alone lack the immune components of the tumour microenvironment required to fully model clinical responses and resistance mechanisms(25,26). Incorporation of immune cells into organoid cultures has enabled evaluation of chemotherapies, targeted therapies, and immunotherapies within more physiologically relevant systems(27–29). Organoid-immune co-culture models have shown promise for predicting immunotherapy responses, informing personalised treatment strategies, and uncovering toxicities or resistance mechanisms that are frequently missed in animal models(30).

Several organoid-immune co-culture methodologies have been reported, including systems based on intact tumour fragments, matrix-embedded patient-derived organoids, autologous immune cells and co-cultures incorporating peripheral blood mononuclear cells (PBMCs) or T cell effectors together with immune engagers(31–34). While these approaches have advanced mechanistic understanding of tumour-immune interactions, matrix composition, culture format, and donor-specific variability can complicate quantitative assessment of T cell motility, cell recovery, and assay-to-assay reproducibility in some screening applications. Moreover, reliance on autologous immune cells introduces practical constraints related to limited cell availability, variable cell quality, and inter-patient variability.

Here, we present a low-matrix suspension organoid-T cell co-culture platform specifically designed for quantitative screening of T cell engagers. By optimizing key assay parameters, we established a standardized suspension co-culture workflow enabling quantitative assessment of tumour killing, immune activation, cytokine secretion, and tumour-immune interactions.

This suspension-based system supports scalable and reproducible evaluation of immune engagers using patient-derived CRC organoids and defined allogeneic T cell populations. Allogeneic CD3+ T cells were incorporated to facilitate standardized evaluation across organoid models while overcoming practical limitations of autologous co-culture systems, including limited cell availability, variable cell quality, and donor-dependent variation. To enable precise effector-to-target ratios and minimize interference from other immune-cell populations, purified CD3+, CD4+, and CD8+ T cell subsets were used. While this approach does not fully recapitulate patient-specific tumour-immune interactions, it enables controlled assessment of immune-engager activity across multiple tumour models and donor backgrounds.

To demonstrate the utility of the platform, we evaluated proof-of-concept T cell engagers targeting HER2 and EGFR. HER2-targeted therapies have shown promising activity in metastatic CRC and are emerging as an important therapeutic strategy in selected patient populations(35). Likewise, EGFR-targeted therapies have been a mainstay of treatment for RAS/BRAF wild-type metastatic CRC for more than a decade, reflecting the frequent expression of EGFR in this disease(36). Using these clinically relevant targets, we sought to establish a human-relevant platform for the functional evaluation of immune-engager activity and the identification of translational biomarkers in CRC.

## METHODS

Experimental procedures, reagent information, culture media compositions, antibody panels, imaging workflows, sequencing protocols, and bioinformatic analyses are provided in the manuscript, **Supplementary Methods** and **Supplementary Table S5**. A detailed protocol is publicly available on **Protocols.io: *A step-by-step low-matrix suspension co-culture method to screen T cell engager activity in patient-derived organoids*** (DOI: https://doi.org/10.17504/protocols.io.kxygxjd1dl8j/v1) (49).

### Patient samples and ethics

CRC specimens were obtained from patients undergoing surgery at University Hospitals Birmingham NHS Foundation Trust, UK, following written informed consent. Ethical approval was granted by the North West-Haydock Research Ethics Committee (REC reference 20/NW/0001; previous REC reference 15/NW/0079; HBRC sub-approval 17-287). All samples were collected and used in accordance with approved ethical protocols and institutional guidelines.

### CRC organoids generation and characterisation

Patient-derived CRC organoids were established from surgical specimens or obtained through the Human Cancer Models Initiative (HCMI). Organoids were cultured under standard conditions(15–17), expanded, cryopreserved, and routinely tested for mycoplasma contamination. Molecular characterization, including microsatellite status, RNA sequencing, whole-genome sequencing, and HLA typing, was performed as described in the **Supplementary Methods**. Clinical and molecular characteristics are provided in **Supplementary Tables S3-S4**.

### T cells isolation and activation

CD3+ T cells were isolated from healthy donor peripheral blood mononuclear cells by immunomagnetic negative selection and expanded following CD3/CD28 stimulation. Donor HLA information is provided in **Supplementary Table S2**.

### Organoid-T-cell co-culture and T cell engager screening

For optimized co-culture assays, CRC organoids were dissociated to single cells and cultured in suspension with 1% BME. Five-day organoids were co-cultured with 7-day activated CD3+ T cells at a 2:1 effector-to-target ratio (20,000:10,000 cells/well) in a 1:1 mixture of organoid and T cell media. EGFR-targeting, HER2-targeting, isotype-control, or vehicle-control T cell engagers were added at the indicated concentrations and cultures were maintained for 72 h without media replacement.

### Flow cytometry

Flow cytometry was used to quantify organoid viability, T cell viability, activation marker expression, and target-antigen expression. Data were acquired using a NovoCyte Penteon flow cytometer and analysed using NovoExpress software. Gating strategies are shown in **Figure 2B**, and antibody details are provided in **Supplementary Table S5**.

### Cytokine analysis

Culture supernatants were analysed using a multiplex cytokine assay to quantify IFN-γ, granzyme B, perforin, and additional cytokines according to the manufacturer’s instructions.

### EGFR and HER2 expression analysis

Cell-surface EGFR and HER2 expression was quantified by antibody binding capacity analysis using flow cytometry. Protein expression and localization were independently validated by multiplex immunofluorescence (mIF) imaging.

### Live-cell imaging

Time-lapse imaging was performed to assess T cell motility and T cell engager-mediated cytotoxicity. Quantitative image analysis was conducted using Gen5 software.

### Genomic and transcriptomic analyses

Whole-genome and transcriptomic sequencing were performed as described in the **Supplementary Methods**. For the WGS analysis raw FASTQ reads were aligned to the GRCh38.13 reference genome using BWA-MEM and duplicates marked using Picard(37,38). SNV/Indel variant calling was performed using MuTect2 using the tumour/normal calling mode; copy number was called using Canvas and structural variants called using Manta(38–40). VCF were annotated using VEP 101(41). RNA sequencing analysis was performed processing raw FASTQ reads with the nf-core/rnaseq pipeline utilising STAR for alignment and Salmon for read counting and all counts were reported in transcripts per million(42).

## QUANTIFICATION AND STATISTICAL ANALYSIS

Statistical analyses and data visualisation were performed using GraphPad Prism v10 (GraphPad Software, San Diego, CA, USA). Data are presented as mean ± standard deviation (SD) unless otherwise stated. Organoid viability, T cell viability, T cell activation, cytokine secretion, motility, and receptor expression data are presented descriptively from independent biological experiments and were not subjected to formal statistical hypothesis testing. Dose-response curves and half-maximal inhibitory concentration (IC50) values were calculated by nonlinear regression using a variable-slope dose– response model in GraphPad Prism. At least three independent biological experiments were performed for all analyses, with technical replicates included where indicated. No samples or data points were excluded from the analyses.

## RESULTS

### Optimization of co-culture medium composition and matrix concentration supports organoid and T cell viability and motility

We investigated the medium and UltiMatrix BME concentration to allow both organoids and CD3+ T cells to grow and expand in co-culture. To identify a co-culture medium that preserved both epithelial and CD3+ T cells viability and most of all did not interfere with the mechanism of action of HER2- and EGFR-directed T cell engagers, we compared seven different media formulations with or without SB202190 (p38-MAPK inhibitor) and Y-27632 (ROCKi: Rho-kinase inhibitor), as both compounds are known to directly modulate T cell activation, proliferation, cytoskeletal organisation and effector function. Specifically, p38-MAPK inhibition enhances T cell proliferation and cytokine production while altering activation thresholds(43), whereas Rho-kinase signalling regulates T cell structural dynamics and clonal expansion(44) and its inhibition can reshape anti-tumour immunity(45). Because HER2- and EGFR-directed T cell engagers rely on intact T cell activation pathways, the presence of p38i or ROCKi could confound cytotoxicity readouts. For this reason, both inhibitors were systematically evaluated and subsequently removed from the final co-culture medium.

Additional organoid-medium components—such as nicotinamide, TGFβ/Smad inhibitors, and other additives—may also influence T cell behaviour, as previously reported(33). However, in this study we specifically tested the medium only after removing p38i and ROCKi, as our optimisation strategy first prioritised components with direct and well-established immunomodulatory effects on T cells. Rather than deconstructing the entire organoid medium at once, we followed a staged, hypothesis-driven approach beginning with factors most likely to influence immune function. This strategy is conceptually aligned with the rational, stepwise refinement of intestinal organoid culture conditions described by Fujii et al.(46) who similarly removed p38i and other inhibitors during medium optimisation to improve physiological relevance. Thus, we tested the following conditions: i. O.M., in-house organoid medium; ii. RPMI + 10% (v/v) FBS; iii. ImmunoCult T cell Expansion Medium; iv. RPMI + 10% (v/v) FBS:OM (80:20 v/v); v. RPMI + 10% (v/v) FBS:O.M.–SB (80:20 v/v), lacking SB202190; vi. OM–SB/Y-27632:ImmunoCult (50:50 v/v), lacking both SB202190 and Y-27632; vii. IntestiCult:ImmunoCult (50:50 v/v) (**Supplementary Table S1**).

We then compared pre-stimulated and unstimulated CD3+ T cells isolated from a healthy donor leukopak (LMX_129) using the different culture media listed above, and co-cultured them with CRC organoids in 100% UltiMatrix BME to assess the media composition effect on organoid growth by brightfield (BF) microscopy (**Fig. 1A**). All isolated CD3+ T cells leukopak donors were assessed for CD25 and CD69 activation marker co-expression at 24, 48, and 72 hours post-isolation before being used in any assay (**Supplementary Fig. S1**).

**Figure 1.**
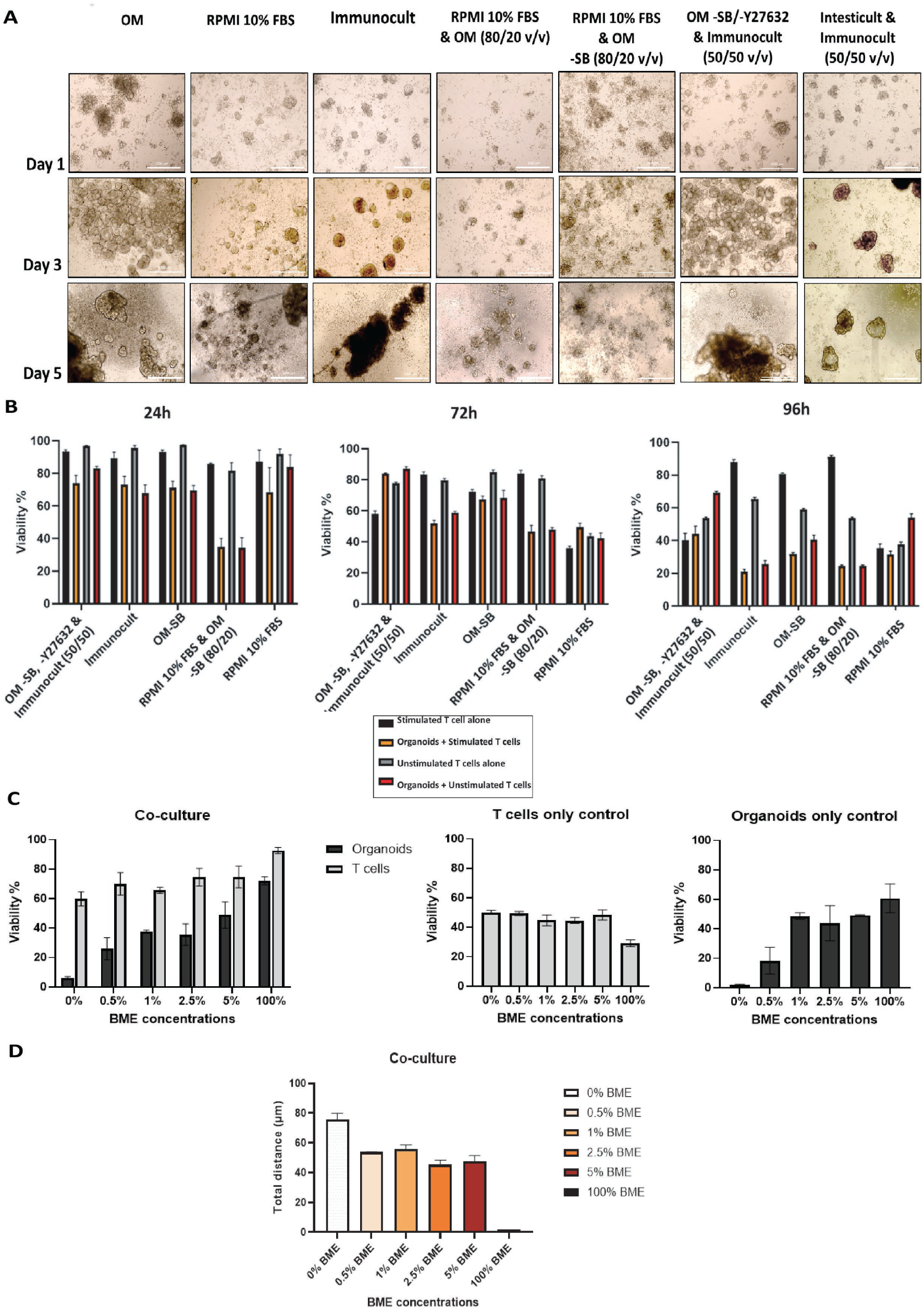
Optimization of organoid-T cell co-culture conditions. **(A)** Representative bright-field images of CRC organoids co-cultured with unstimulated CD3+ T cells under the indicated culture conditions (**Supplementary Table S1**). Images were acquired at 10× magnification. Scale bars, 1000 μm. **(B)** Viability of organoids and stimulated or unstimulated CD3+ T cells after 24, 72, and 96 h of co-culture. **(C-D)** Viability and motility of stimulated CD3+ T cells following 72 h of co-culture under increasing UltiMatrix BME concentrations. Viability was assessed by flow cytometry and normalised to the corresponding organoid monoculture control at the same time point. Additional co-culture controls and T cell activation data are shown in **Supplementary Fig. S2.** Data represent mean ± SD from at least three independent biological experiments, each performed with technical replicates.

We assessed organoids and CD3+ T cells viability in co-culture and controls at 24, 72 and 96 hours (**Fig. 1B**) as well as CD3+ T cells activation in co-culture by flow cytometry (**Supplementary Fig. S2**).

Across all conditions, the medium that best supported both organoid integrity and T cell viability— particularly beyond 72 hours was the in-house organoid medium lacking SB202190 and Y-27632, diluted 1:1 with ImmunoCult (v/v). Importantly, removal of p38i and ROCKi did not impair organoid growth in co-culture, confirming that these inhibitors are dispensable for organoids culture under these conditions but detrimental to accurate assessment of T cell engager activity.

Next, we focused on improving CD3+ T cells viability and mobility thus killing capacity in co-culture. To this end, we evaluated the effect of basement membrane extract (UltiMatrix BME) concentration on T cell behaviour using UltiMatrix BME diluted to 0%, 0.5%, 1%, 2.5%, 5% or 100% in a 50:50 (v/v) suspension mixture of OM–SB/Y-27632:ImmunoCult. CRC organoids were cultured for 5 days under the indicated UltiMatrix BME conditions and subsequently co-cultured for 72 hours with 7-day pre-stimulated CD3+ T cells (donor 054) stained with CellTracker Deep Red dye. Co-cultures were maintained in a BioSpa 8 Automated Incubator, and time-lapse imaging was performed every 12 hours using a BioTek Cytation 5 Cell Imaging Multi-Mode Reader with bright-field and Cy5 channels at 10x magnifications (**Supplementary Videos S1A–F**). To quantitatively assess CD3+ T cells-motility, image acquisition and time-lapse sequences were analysed using Gen5 software by tracking individual CD3+ T cells over the 72 hours imaging period, and total track distance was extracted for each condition. These analyses revealed a progressive reduction in CD3+ T cells-motility with increasing UltiMatrix BME concentration. In particular, in 100% UltiMatrix BME CD3+ T cells deposit by gravity at the bottom of the well significantly reducing CD3+ T cells movement compared with 1% (v/v) UltiMatrix BME, consistent with a marked decrease in total track distance (**Supplementary Videos S1A–F**). In parallel, CD3+ T cell viability was quantified by flow cytometry (Novocyte Panteon, Agilent) and showed that although 100% UltiMatrix BME supported high overall viability in fully embedded co-cultures, low UltiMatrix BME concentrations ( < 1% (v/v)) negatively affected organoids growth, while high UltiMatrix BME concentrations impaired CD3+ T cells movement and effectively restricted CD3+ T cells movement (**Fig. 1C–D; Supplementary Videos S1A–F**).

Based on these quantitative assessments and subsequent co-culture screening experiments, 1% (v/v) UltiMatrix BME in OM-SB/Y-27632:ImmunoCult (50:50, v/v) was selected as the optimal condition, providing a balance between CD3+ T cell viability, motility, and preservation of organoid structural integrity while maximising organoid killing efficiency. Around day 5, CRC organoids transition from an early fetal-like/regenerative transcriptional state to a more mature cancer-like epithelial phenotype(16,47,48). Using organoids at this stage improves the physiological relevance of drug-response assays by better reflecting tumour-specific pathways, differentiation states, and cellular heterogeneity(30). Accordingly, and considering the 72-hour duration of the T-cell engager (TCE) assay, organoids were screened 5 days after single-cell seeding. These findings informed the final suspension co-culture workflow, in which 7-day-activated CD3+ T cells and 5-day-mature CRC organoids were co-cultured with TCEs for 72 hours in 1% (v/v) UltiMatrix BME diluted in OM-SB/Y-27632:ImmunoCult (50:50, v/v), without media exchange throughout the assay period (**Fig. 2A**). The optimised workflow, together with the flow-cytometry gating strategy shown in **Fig. 2B**, was used for all subsequent proof-of-concept studies.

**Figure 2.**
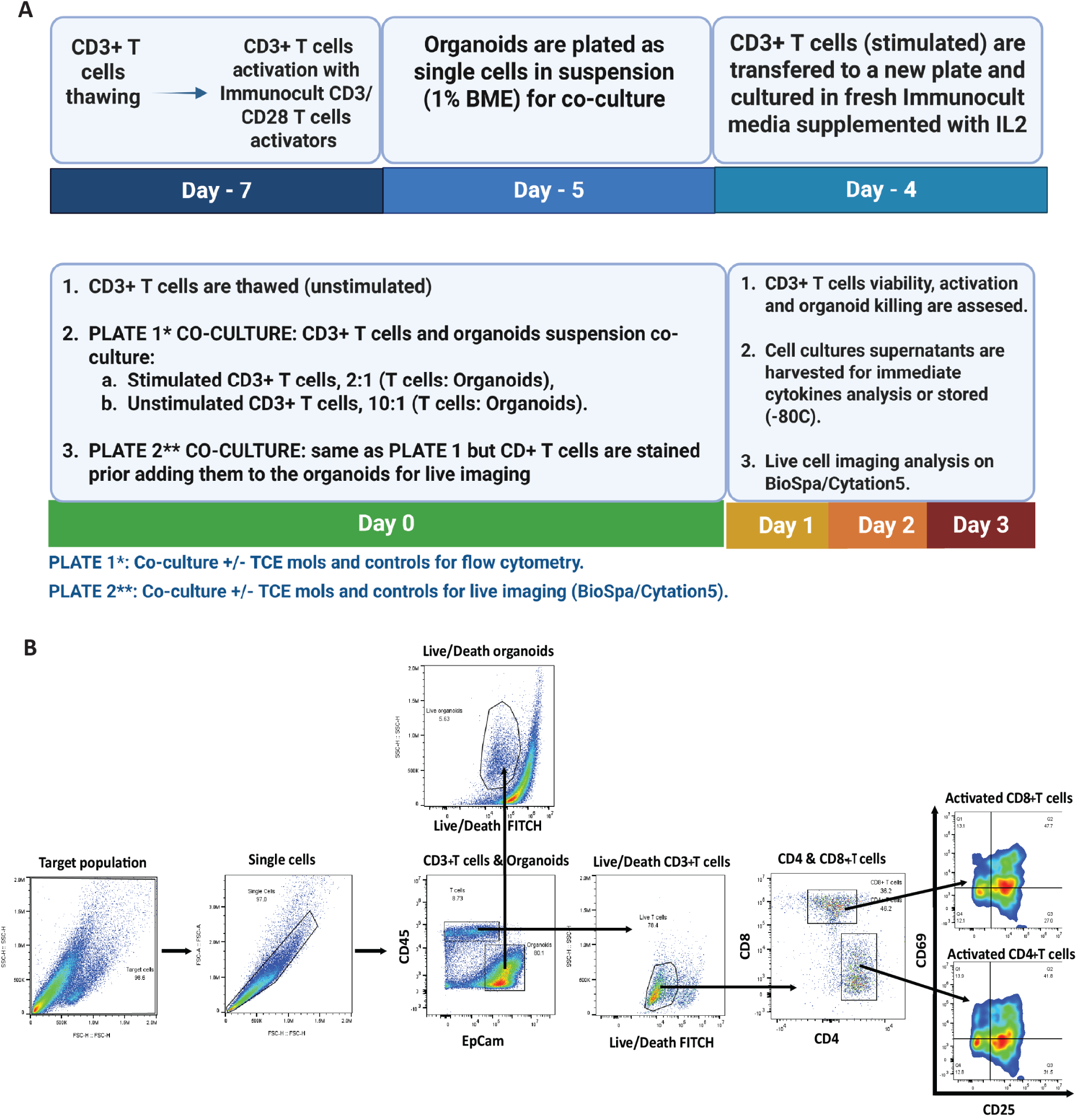
Immune engagers screening protocol and gating strategy. **(A)** Suspension co-culture method for screening immuno-oncology compounds using CD3+ T cells and CRC organoids models. Created with BioRender.com **(B)** Sequential flow cytometry gating strategy for singlets, organoids viability and CD3, CD4 and CD8+ T cells viability and activation post co-culture. The gating strategy is shown by the black arrows. Compensation of data was performed in the NovoExpress software.

### EGFR-TCE induces reproducible organoid killing and T cell activation across donors

Having established the optimal co-culture workflow, we next evaluated the assay duration required to assess T cell activation and tumour organoid killing. CRC organoids (COLO250, COLO298, and S315571) were co-cultured with 7-day-activated CD3+ T cells from four healthy donors (105C, 804C, 702C, and LMX_129; **Supplementary Table S2**) and treated with EGFR-TCE (100 nM) or PBS vehicle control for 48 or 72 hours (Fig. 3A-B). Increased EGFR-TCE-mediated killing was observed at 72 h compared with 48 h, with average increases of 3.97% (105C), 21.35% (804C), 19.84% (702C), and 13.06% (LMX_129) (**Fig. 3A**). CD3+ T-cell viability remained above 50% across all treatment conditions (**Fig. 3B**). Donor 105C exhibited the greatest increase in CD4+ and CD8+ T cell activation following EGFR-TCE treatment (**Supplementary Fig. S3**), which was associated with higher IFN-γ levels in the corresponding co-cultures (**Fig. 3C**). Nevertheless, activation and IFN-γ production were observed across all donor-derived CD3+ T cell co-cultures following EGFR-TCE treatment. Matched control conditions are shown in **Supplementary Fig. S4**.

**Figure 3.**
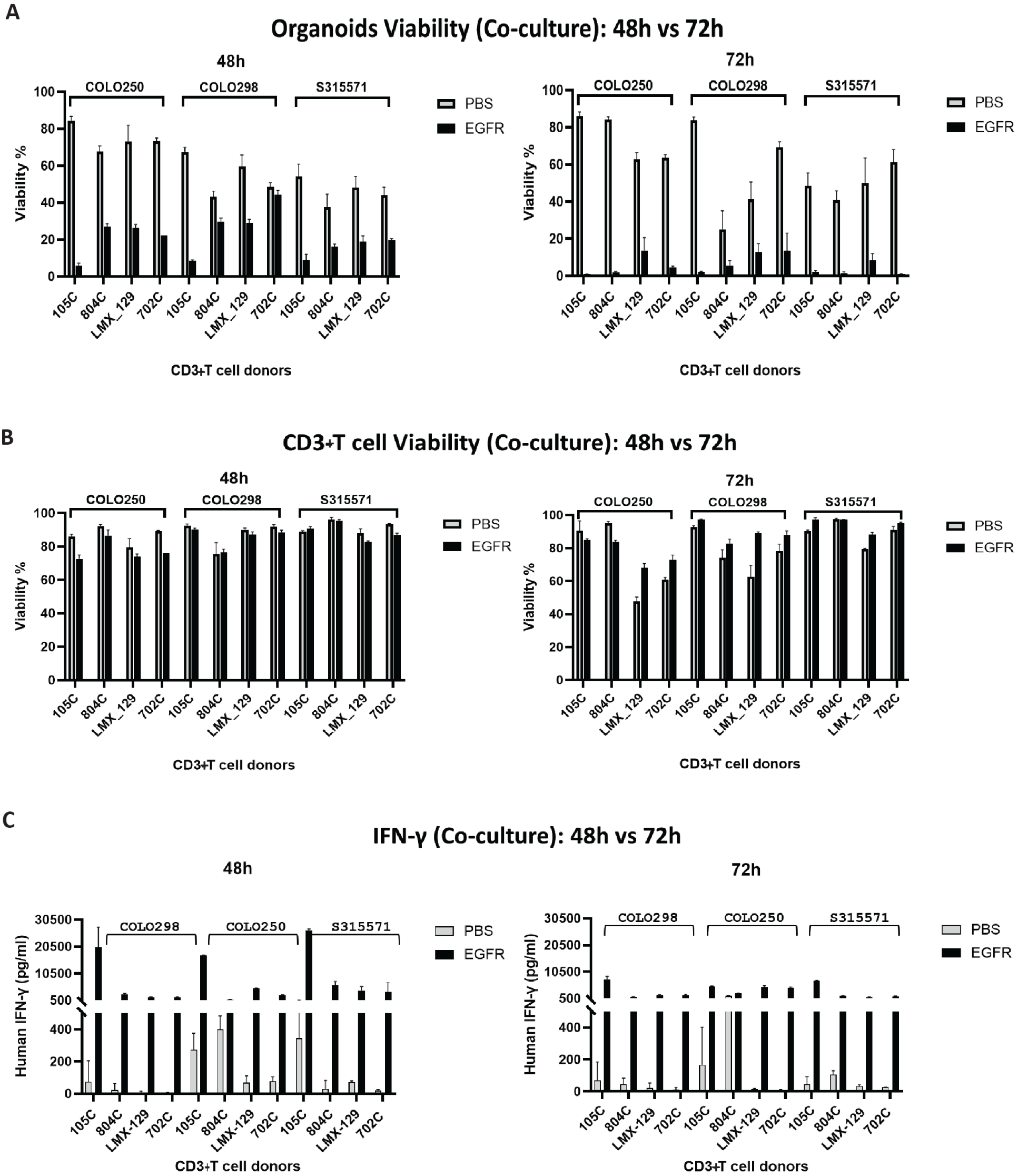
EGFR-TCE-mediated effects in CRC organoid-T cell co-cultures. CRC organoids (COLO250, COLO298, and S315571) were co-cultured with stimulated CD3+ T cells from healthy donors (105C, 804C, 702C, and LMX_129) and treated with EGFR-TCE (100 nM) or PBS for 48 and 72 h. **(A)** Organoid viability. Viability was measured via flow cytometry viability staining. Values were normalized to the respective organoid model as mono-culture at the indicated time point. **(B)** CD3+ T cell viability. **(C)** IFN-γ secretion in co-culture supernatants. Matched control conditions and T cell activation data are shown in **Supplementary Fig. S3 and S4.** Data represent mean ± SD from at least three independent biological experiments, each performed with technical replicates.

### CRC organoid screening identifies models suitable for immune engager profiling

We next sought to identify CRC organoid-T cell co-culture models suitable for immune engager screening, defined by low non-specific T cell cytotoxicity and baseline organoid viability ≥50% in PBS-treated co-cultures. Twenty-one colorectal tumour organoid models representing colon and rectal cancers, MSS and MSI subtypes, CMS1-4 classifications, diverse genomic backgrounds, different HLA antigen expression profiles, and variable EGFR and HER2 expression levels were screened (**Supplementary Tables S3-S4**). CRC organoids were co-cultured with stimulated CD3+ T cells from donor 105C at a 2:1 effector-to-target ratio and treated with EGFR-TCE (100 nM) or PBS for 72 h. Organoid viability, CD3+ T cell viability, and CD4+ and CD8+ T cell activation (CD69 and CD25) were assessed by flow cytometry (**Fig. 4; Supplementary Fig. S5**).

**Figure 4.**
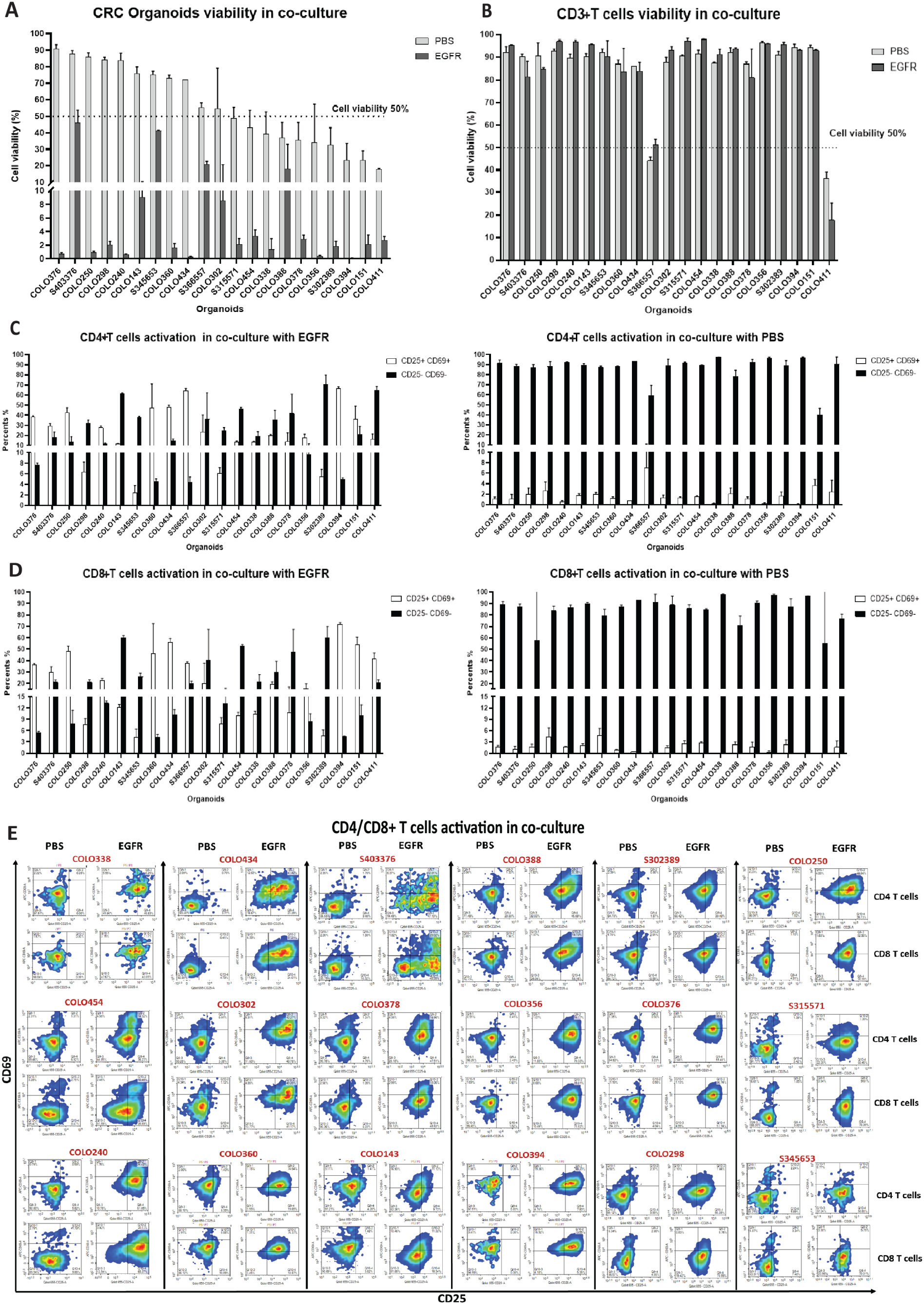
CRC organoid and stimulated CD3+ T cell (105C donor) co-culture viability screening following treatment with EGFR-TCE (100 nM) or PBS at 72 h. **(A)** Viability of 21 CRC organoid models. Viability was assessed via flow cytometry viability staining and normalized to the respective organoid model as mono-culture at the indicated time point. **(B)** Viability of stimulated CD3+ T cells. **(C)** CD25 and CD69 expression in stimulated CD4+ T cells. **(D)** CD25 and CD69 expression in stimulated CD8+ T cells. **(E)** Activation of stimulated CD4+ and CD8+ T cells. Matched control conditions are shown in **Supplementary Fig. S5**. Data represent mean ± SD from at least three independent biological experiments, each performed with technical replicates.

Organoid viability in PBS-treated co-cultures remained ≥50% in 12 of 21 models, whereas stimulated CD3+ T cell viability exceeded 80% in 19 models irrespective of treatment (**Fig. 4A-B**). EGFR-TCE treatment increased CD25 and CD69 expression in both CD4+ and CD8+ T cells relative to PBS controls (**Fig. 4C-E**). Of the 12 models meeting the viability threshold, COLO298, COLO250, and S345653 were selected for subsequent EGFR-TCE and HER2-TCE dose-response studies based on their robust co-culture performance and distinct target-expression profiles. Notably, S345653 displayed partial sensitivity to EGFR-TCE, with approximately 40% loss of viability at 100 nM, whereas COLO298 and COLO250 exhibited near-complete loss of viability under the same conditions (**Fig. 4A**).

### Target expression in selected CRC organoid models

COLO298, COLO250, and S345653 exhibited distinct EGFR and HER2 expression profiles (**Supplementary Table S3**) and were subsequently used to evaluate antigen-dependent responses to EGFR- and HER2-targeting T cell engagers. The models also exhibited diverse morphological features, ranging from simple cystic structures with large lumens to densely packed organoids with smaller lumens, which may influence T cell infiltration and cytotoxicity (**Fig. 5**).

**Figure 5.**
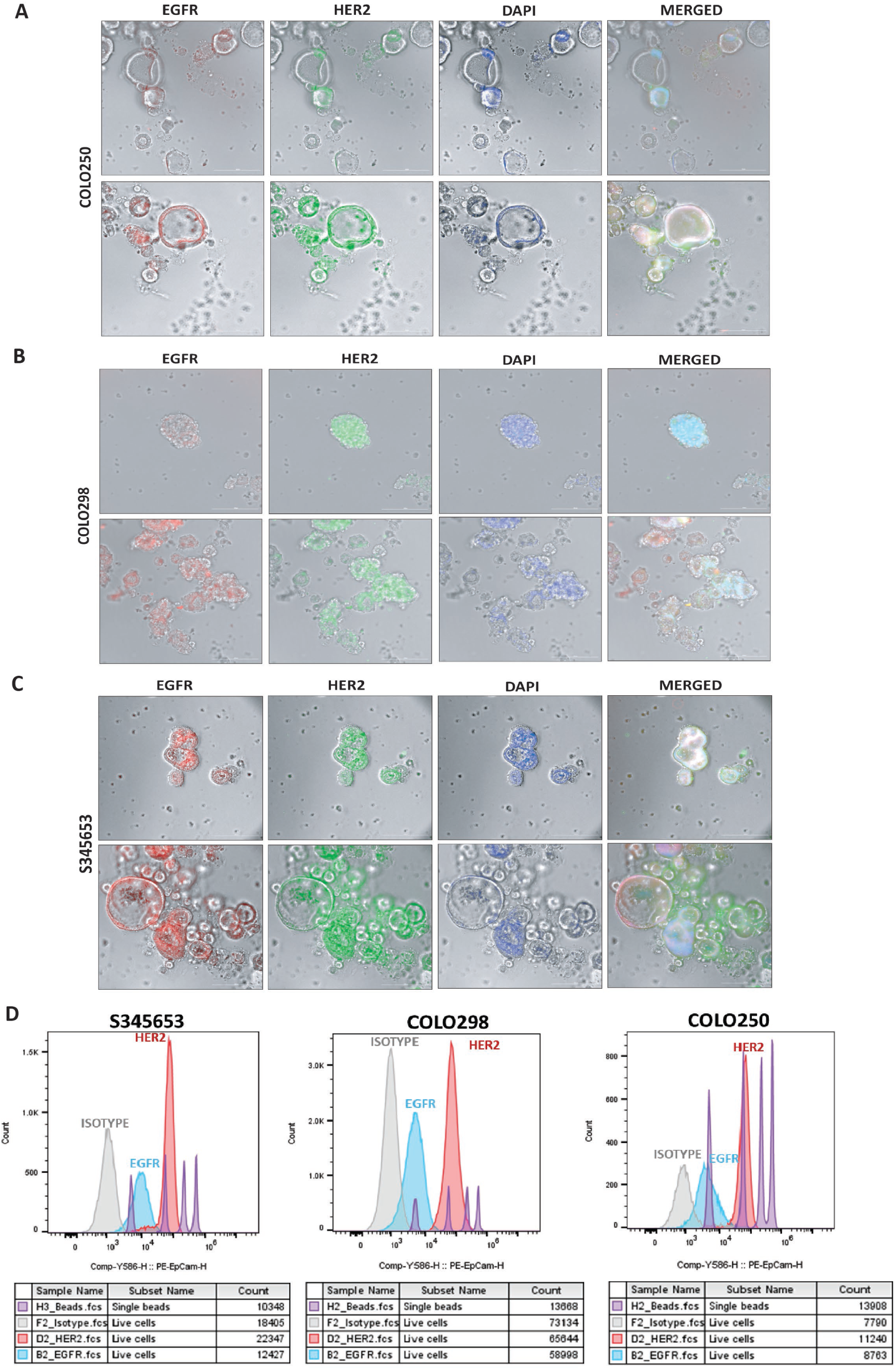
EGFR and HER2 expression in patient-derived colorectal cancer organoids. **(A-C)** Representative multiplex immunofluorescence images of day-5 COLO250, COLO298, and S345653 organoids showing EGFR, HER2, and DAPI staining. Bright-field overlays and merged images are shown. Scale bars, 200 μm. **(D)** Quantification of cell-surface EGFR and HER2 expression in COLO250, COLO298, and S345653 organoids by flow cytometry. Receptor expression is presented as antibody binding capacity (ABC) values. Data represent mean ± SD from at least three independent biological experiments, each performed with technical replicates.

Multiplex immunofluorescence (mIF) analysis confirmed the presence and membrane localisation of EGFR and HER2 protein targets in all three CRC organoid models selected for the screening (**Fig. 5A–C**), supporting their suitability for evaluating target-dependent TCE activity.

We also analysed EGFR and HER2 surface receptor expression using Quantibrite PE Phycoerythrin Fluorescence Quantitation beads in COLO298, COLO250, and S345653 organoid models by flow cytometry, enabling conversion of PE fluorescence into antibody binding capacity (ABC) values. Across all three models, HER2 surface expression was consistently higher than EGFR (**Fig. 5D**).

### HER2- and EGFR-targeting TCEs demonstrate distinct dose-response activity

We next evaluated the activity of HER2-TCE and EGFR-TCE in COLO298, COLO250, and S345653 organoid-T cell co-cultures. Organoids were treated with increasing concentrations of HER2-TCE, EGFR-TCE, or isotype-TCE for 72 h, and organoid viability, CD3+ T cells viability, and T cells activation were assessed by flow cytometry (**Fig. 6A-B; Supplementary Fig. S6**).

**Figure 6.**
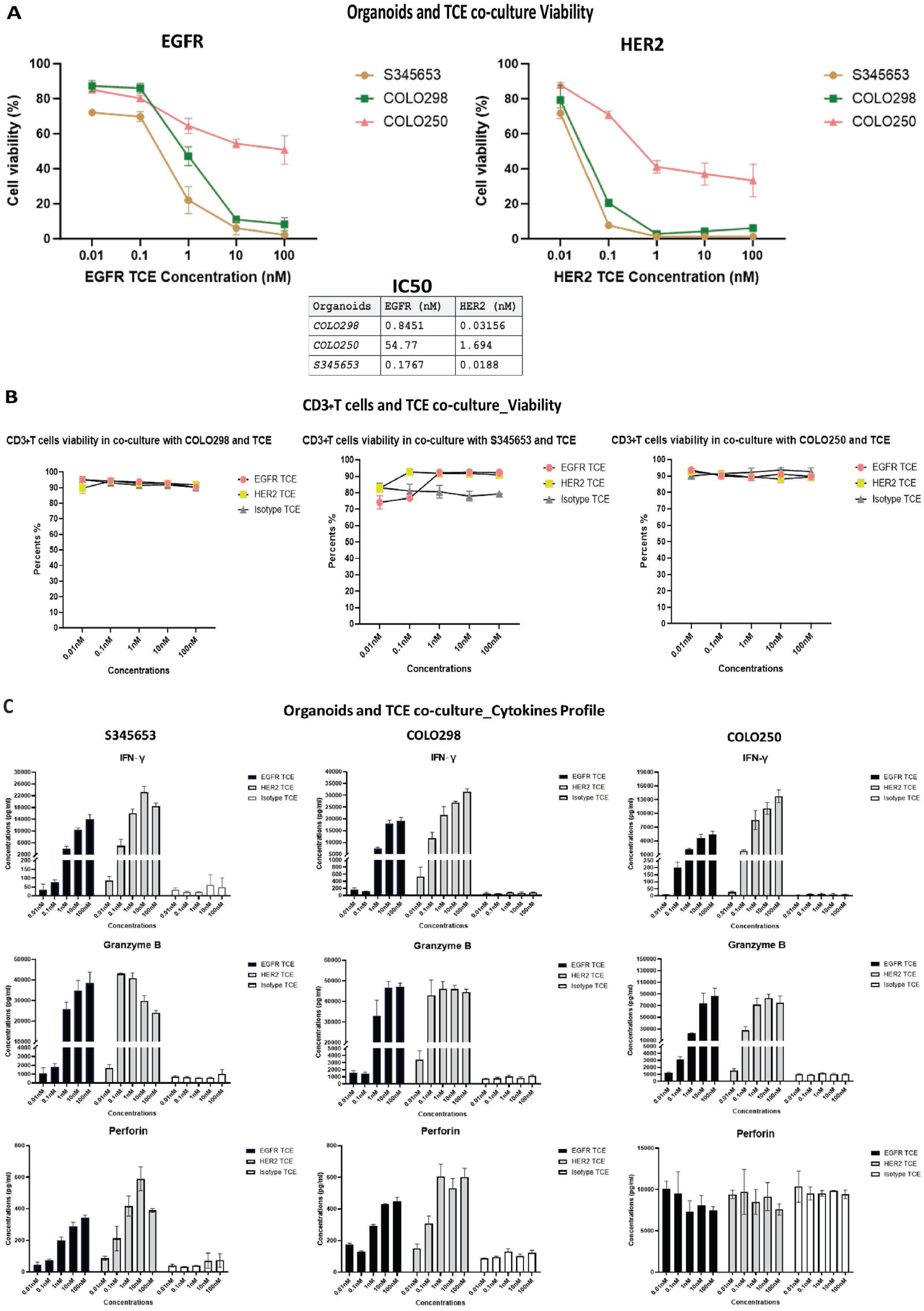
Functional activity of T-cell engagers in CRC organoid-T cell co-cultures. CRC organoids (S345653, COLO298 and COLO250) were co-cultured with stimulated CD3+ T cells and treated with increasing concentrations of HER2-TCE, EGFR-TCE or isotype-TCE for 72 h. **(A)** Dose-response curves showing organoid viability and corresponding IC₅₀ values. **(B)** CD3+ T cell viability following co-culture and TCE treatment. **(C)** IFN-γ, granzyme B and perforin concentrations in co-culture supernatants after 72 h. Additional T cell activation and control data are shown in **Supplementary Fig. S6 and S7.** Data represent mean ± SD from three independent biological experiments, each performed with technical replicates. IC_50_ values were calculated by nonlinear regression using a variable-slope dose–response model in GraphPad Prism.

S345653 and COLO298 organoids were highly sensitive to both TCEs, with lower IC₅₀ values than COLO250. For S345653, IC₅₀ values were 0.0188 nM for HER2-TCE and 0.1767 nM for EGFR-TCE, while COLO298 showed IC₅₀ values of 0.0316 nM and 0.8451 nM, respectively. In contrast, COLO250 organoids exhibited substantially reduced sensitivity, with IC₅₀ values of 1.694 nM for HER2-TCE and 54.77 nM for EGFR-TCE (**Fig. 6A**). Consistent with these values, HER2-TCE induced marked cytotoxicity in S345653 and COLO298 organoids at concentrations as low as 0.1 nM, whereas EGFR-TCE activity became apparent from 1 nM onwards. COLO250 organoids displayed a comparatively attenuated response to both TCEs (**Fig. 6A**).

Time-lapse imaging confirmed TCE-mediated tumour organoid killing, revealing progressive caspase-3/7 activation and accumulation of CD3+ T cells around organoids in a concentration-dependent manner following treatment with either HER2-TCE or EGFR-TCE. Minimal apoptotic activity was observed in organoid-only controls treated with TCE molecules or in T cell-only controls (**Supplementary Videos S2-4A-J**).

To further characterise T cell function, IFN-γ, granzyme B, and perforin secretion were quantified in co-culture supernatants after 72 h of treatment (**Fig. 6C**). Both HER2-TCE and EGFR-TCE induced robust, dose-dependent cytokine production across the organoid models tested. In contrast, isotype-TCE controls did not induce detectable IFN-γ, granzyme B, or perforin secretion. Cytokine production was also detected in T cell-only cultures exposed to TCEs, confirming direct T cell activation by the engager molecules (**Supplementary Fig. S7**). Additional cytokines, including TNF-α, IL-10, IL-6, IL-2, and IL-17A, were not detected under the conditions tested.

A discrepancy remains between the EGFR-TCE cytotoxicity observed in **Fig. 4** and **Fig. 6** for S345653 and COLO250 organoids. The screening results in **Fig. 4** and **Supplementary Fig. S5** were generated using earlier-passage (P ≤ 20) S345653, COLO298, and COLO250 organoids, whereas the data in **Fig. 5–6** and **Supplementary Fig. S6–7** were obtained from later-passage (P ≤ 30) cultures. As the experiments were performed using different organoid passages, these findings suggest that although TCE-mediated killing is reproducible across co-culture screens, the magnitude of the response may vary with organoid passage number.

## DISCUSSION

Colorectal cancer (CRC) remains a leading cause of cancer-related mortality worldwide, and despite recent therapeutic advances, treatment is frequently limited by toxicity, resistance, and disease recurrence(1). In this context, the development of human-relevant *in vitro* platforms capable of modelling immune-mediated tumour killing and predicting therapeutic responses represents an important component of preclinical immuno-oncology research.

Organoid–immune cell co-culture systems have emerged as powerful tools to interrogate tumour–immune interactions, including approaches combining CRC patient-derived organoids with peripheral blood mononuclear cells or purified T cell populations in the presence of immune engagers(31–34). While these systems have yielded valuable mechanistic insights, their broader application in drug discovery and screening workflows has been constrained by experimental variability introduced by matrix-embedded culture formats, complex or immunomodulatory media components, and limited capacity for quantitative analysis of parameters such as T cell motility, activation, and viability. These challenges complicate standardisation across donors and organoid models and hinder reproducibility at scale.

In this study, we describe a low-matrix suspension co-culture method designed to support systematic and reproducible evaluation of T cell engager activity using patient-derived CRC organoids (PDOs) and allogeneic CD3+ T cells. The suspension co-culture screening protocol provides a robust *in vitro* framework for assessing immune engager activity using CRC PDO models. Following organoid establishment, expansion, quality control, and biobanking, models could be recovered from frozen stocks and deployed in immune-engager co-culture assays within approximately two weeks, assuming prior validation of organoid models.

The method builds on existing organoid–immune co-culture approaches by systematically optimising experimental variables known to influence assay performance, including extracellular matrix concentration, culture medium composition, T cell activation state, and co-culture duration. Adoption of a low-matrix suspension format enables quantitative assessment of T cell motility, viability, activation, and cytotoxic function while preserving key three-dimensional features of organoid architecture. This configuration supports reproducible analysis across multiple organoid genotypes and T cell donors and remains compatible with medium-throughput screening workflows, addressing practical limitations associated with matrix-embedded cultures and autologous immune-cell dependence. The use of defined allogeneic CD3+ T cell populations enabled standardised and scalable screening across organoid models and donors, facilitating comparative assessment of T cell engager activity while reducing the impact of limited availability and functional heterogeneity of autologous patient-derived immune cells.

Application of this platform revealed heterogeneous responses to EGFR- and HER2-targeting T cell engagers across CRC organoid models, consistent with inter-patient biological variability observed clinically. Across models, susceptibility to immune-engager-mediated cytotoxicity was most closely associated with target antigen expression, highlighting target abundance as a potential predictive biomarker of sensitivity to T cell engager therapy. The ability to capture multiple functional endpoints— including organoid killing, caspase-3/7-dependent apoptosis, T cell activation, and secretion of effector cytokines such as IFN-γ, granzyme B, and perforin—supports the utility of this system for integrated functional profiling of immune engager activity. These features support the use of the platform for preclinical candidate prioritisation and biomarker discovery, while providing a functional framework for investigating mechanisms of sensitivity and resistance to immune-engaging therapies.

The clinical relevance of PDO-based platforms is supported by studies demonstrating correlations between organoid responses, treatment sensitivity, and patient outcomes(18,28).

Building on these observations, incorporation of immune components into PDO systems may further enhance the ability to model therapeutic response and resistance to immunotherapies and other immune-engaging therapeutics. Although the study focuses on CRC-PDOs, the protocol can be adapted to other PDO models, including organoids derived from additional tumour types, with co-culture duration adjusted according to the growth characteristics of individual organoid models.

Together, these findings demonstrate that controlled optimisation of co-culture conditions can substantially improve the reproducibility, interpretability, and scalability of organoid-based immune engager assays. Future studies integrating organoid-T cell co-culture responses with clinical outcome data from patients treated with immune-engaging therapies will be required to define the predictive value of this platform for patient stratification and biomarker-guided therapeutic selection, while supporting efforts to reduce reliance on animal models during preclinical immunotherapy development.

### Limitations of the study

Although the suspension co-culture system enables controlled analysis of tumour organoid-T cell interactions, it does not fully recapitulate the complexity of the *in vivo* tumour microenvironment, including stromal, vascular, and immunosuppressive components. The use of allogeneic T cells does not capture patient-specific immune features such as antigen specificity, exhaustion, and dysfunction that may influence clinical responses to T cell engager therapies, while the use of autologous immune cells remains limited by cell availability and donor-to-donor variability. In addition, the suspension format is less suited than matrix-embedded systems for studying immune-cell infiltration and long-term spatial interactions. Future incorporation of stromal and myeloid populations, physiologically relevant microenvironmental conditions, and autologous immune cells may further enhance the translational relevance of the platform. Despite these limitations, the assay provides a robust, scalable, and human-relevant framework for evaluating T cell engager activity, enabling integrated assessment of tumour killing, T cell activation, cytokine secretion, and tumour-immune interactions in patient-derived colorectal cancer models, with potential applications in translational immunotherapy research, candidate prioritisation, biomarker discovery, and patient stratification.

## Supporting information

Videos S1A_F_BME concentration effects on CD3+ T cells movement

Videos S2-4A-J_EGFR HER2 engager cytotoxicity in co-culture

## LIST OF ABBREVIATIONS

ABC: Antibody Binding Capacity
ASC: Adult Stem Cell
BME: Basement Membrane Extract
bsAb: Bispecific Antibody
CMS: Consensus Molecular Subtype
CRC: Colorectal Cancer
DAPI: 4′,6-Diamidino-2-Phenylindole
DMEM: Dulbecco’s Modified Eagle Medium
E:T: Effector-to-Target Ratio
EGF: Epidermal Growth Factor
EGFR: Epidermal Growth Factor Receptor
EGFR-TCE: Epidermal Growth Factor Receptor-Targeting T cell Engager
EMVI: Extramural Vascular Invasion
EpCAM: Epithelial Cell Adhesion Molecule
FACS: Fluorescence-Activated Cell Sorting
FBS: Fetal Bovine Serum
FDA: Food and Drug Administration
HCMI: Human Cancer Models Initiative
HER2: Human Epidermal Growth Factor Receptor 2
HER2-TCE: HER2-Targeting T cell Engager
HLA: Human Leukocyte Antigen
IC50: Half-Maximal Inhibitory Concentration
IFN-γ: Interferon Gamma
IL-2: Interleukin-2
mIF: Multiplex Immunofluorescence
MSI: Microsatellite Instability
MSI-H: Microsatellite Instability-High (if used in text)
MSS: Microsatellite Stable
MTA: Material Transfer Agreement
NK: Natural Killer
O.M.: Organoid Medium
PBMC: Peripheral Blood Mononuclear Cell
PBS: Phosphate-Buffered Saline
PDO: Patient-Derived Organoid
PGE2: Prostaglandin E2
PNI: Perineural Invasion
POC: Proof-of-Concept
POC-TCE: Proof-of-Concept T cell Engager
R&D: Research and Development
RNA-seq: RNA Sequencing
ROCK: Rho-Associated Coiled-Coil Containing Protein Kinase
SD: Standard Deviation
TCE: T cell Engager
TMB: Tumour Mutational Burden
TME: Tumour Microenvironment
TNF-α: Tumour Necrosis Factor Alpha
TPM: Transcripts Per Million
WGS: Whole-Genome Sequencing

## Abbreviations used in Supplementary Table S3

LCCRT: Long-Course Chemoradiotherapy
SCRT: Short-Course Radiotherapy
FOLFOX: Folinic Acid, Fluorouracil and Oxaliplatin Chemotherapy Regimen
LVI: Lymphovascular Invasion

## DECLARATIONS

### Availability of data and material

Organoid models are available from the corresponding author upon reasonable request and completion of a Material Transfer Agreement, subject to applicable ethical approvals. A subset of models is available through the Human Cancer Models Initiative (HCMI) collection at ATCC.

Sequencing datasets have been deposited in the Cell Model Passports database (https://cellmodelpassports.sanger.ac.uk/) and in the NCBI Sequence Read Archive (SRA) under BioProject accession PRJNA978372 (https://www.ncbi.nlm.nih.gov/bioproject/PRJNA978372).

### Ethics approval and consent to participate

Informed consent was obtained from all participants. Pseudonymised clinical data in this study include patients’ gender.

### Authors’ contributions

**Conceptualization:** C.M.A.P., A.D.B., S.J.D.

**Methodology:** C.M.A.P., J.Z., K.R., L.T., D.G.M., C.B., N.L., E.R.

**Investigation:** C.M.A.P., J.Z., K.R., L.T., L.B., K.A.M.

**Resources:** D.G.M., C.B., E.R., O.H., M.G., R.L.

**Data interpretation:** C.M.A.P., J.Z., D.G.M., C.B.

**Funding acquisition:** A.D.B.

**Supervision:** C.M.A.P., A.D.B., S.J.D., S.C.

**Writing – original draft:** C.M.A.P.

**Writing – review & editing:** C.M.A.P., J.Z., A.D.B., D.G.M., C.B., S.J.D.

**All authors reviewed and approved the final manuscript.**

### Lead contact

Requests for further information and resources should be directed to and will be fulfilled by the corresponding author, Claudia M. A. Pinna.

## Acknowledgements

The authors extend their sincere thanks to the patients who participated in this study and contributed samples. We gratefully acknowledge the support of the University of Birmingham’s Human Biomaterials Resource Centre, which was originally established through the Birmingham Science City – Experimental Medicine Network of Excellence Project.

The authors also thank Dr Amanda Barnes (Agilent) for her technical guidance and support with the live-imaging assay setup and analysis.

**Figure 2A** and the graphical abstract were created in BioRender.

## SUPPLEMENTARY INFORMATION

**Supplementary Document S1. Supplementary Methods, Fig. S1-S7, and Tables S1-S5.**

**Table S5.** Key Resources Table listing biological materials, antibodies, reagents, assays, software, and instrumentation used in this study.

**Videos S1A-S1F.** Effect of basement membrane extract (BME) concentration on CD3+ T cell motility in CRC organoid-T cell co-cultures (related to **Fig. 1).**

**Videos S2A-S4J.** T cell engager-mediated cytotoxicity in CRC organoid-T cell co-cultures and matched control conditions (related to **Fig. 6).**

---

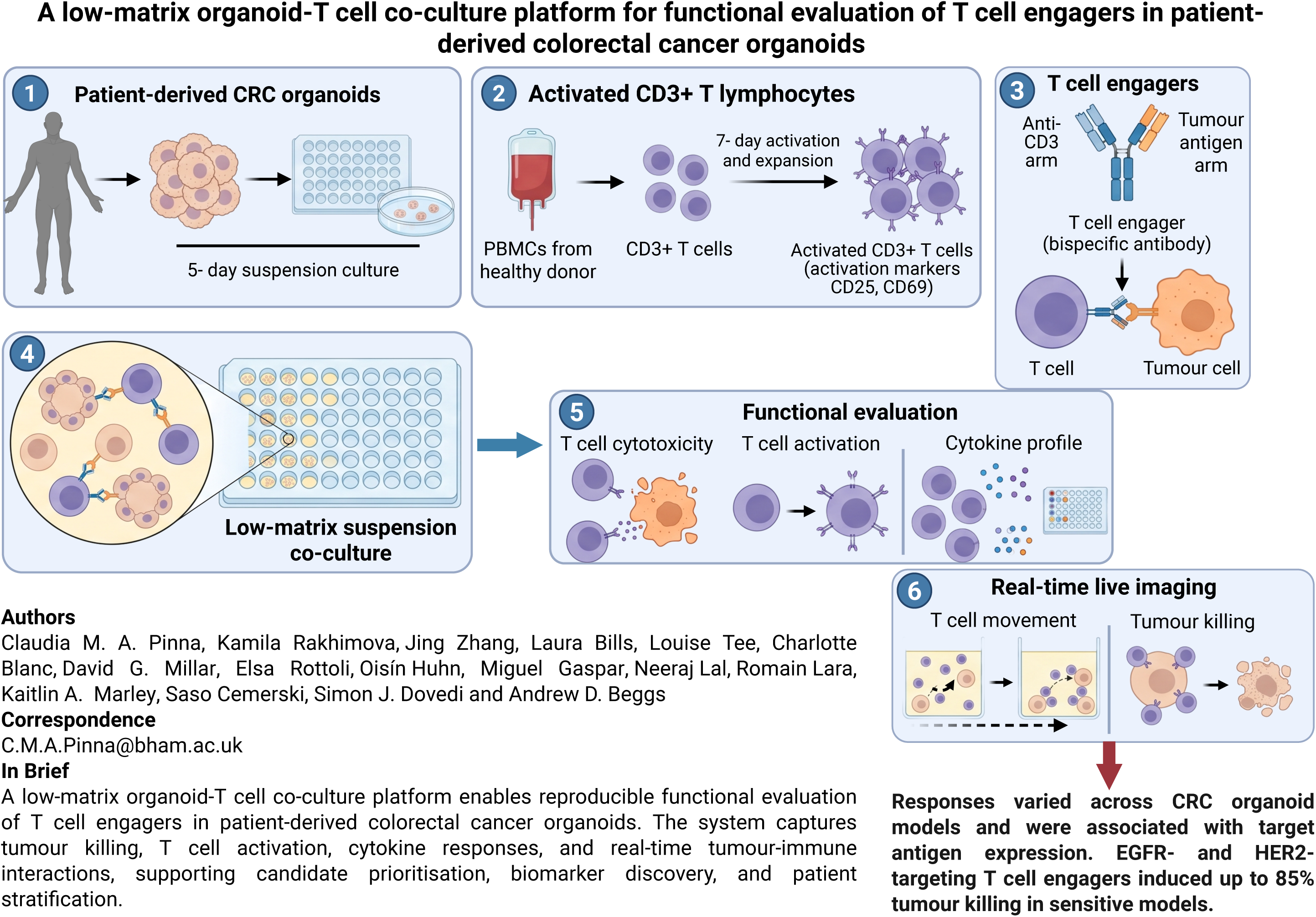

## REFERENCES

1. Ferlay J, Soerjomataram I, Dikshit R, Eser S, Mathers C, Rebelo M, et al. Cancer incidence and mortality worldwide: sources, methods and major patterns in GLOBOCAN 2012. Int J Cancer. 2015 Mar 1;136(5):E359–386. doi:10.1002/ijc.29210 PubMed PMID: 25220842.

2. Van Cutsem E, Cervantes A, Adam R, Sobrero A, Van Krieken JH, Aderka D, et al. ESMO consensus guidelines for the management of patients with metastatic colorectal cancer. Ann Oncol. 2016 Aug;27(8):1386–422. doi:10.1093/annonc/mdw235 PubMed PMID: 27380959.

3. Junttila MR, de Sauvage FJ. Influence of tumour micro-environment heterogeneity on therapeutic response. Nature. 2013 Sep 19;501(7467):346–54. doi:10.1038/nature12626 PubMed PMID: 24048067.

4. Galon J, Mlecnik B, Bindea G, Angell HK, Berger A, Lagorce C, et al. Towards the introduction of the ‘Immunoscore’ in the classification of malignant tumours. The Journal of Pathology. 2014;232(2):199–209. doi:10.1002/path.4287

5. Thakur A, Huang M, Lum LG. Bispecific antibody based therapeutics: Strengths and challenges. Blood Rev. 2018 Jul;32(4):339–47. doi:10.1016/j.blre.2018.02.004 PubMed PMID: 29482895.

6. Fenis A, Demaria O, Gauthier L, Vivier E, Narni-Mancinelli E. New immune cell engagers for cancer immunotherapy. Nat Rev Immunol. 2024 Jan 25. doi:10.1038/s41577-023-00982-7 PubMed PMID: 38273127.

7. Li H, Er Saw P, Song E. Challenges and strategies for next-generation bispecific antibody-based antitumor therapeutics. Cell Mol Immunol. 2020 May;17(5):451–61. doi:10.1038/s41423-020-0417-8 PubMed PMID: 32313210; PubMed Central PMCID: PMC7193592.

8. Wong CH, Siah KW, Lo AW. Estimation of clinical trial success rates and related parameters. Biostatistics. 2019 Apr;20(2):273–86. doi:10.1093/biostatistics/kxx069 PubMed PMID: 29394327; PubMed Central PMCID: PMC6409418.

9. Voskoglou-Nomikos T, Pater JL, Seymour L. Clinical predictive value of the in vitro cell line, human xenograft, and mouse allograft preclinical cancer models. Clin Cancer Res. 2003 Sep 15;9(11):4227–39. PubMed PMID: 14519650.

10. Mak IW, Evaniew N, Ghert M. Lost in translation: animal models and clinical trials in cancer treatment. Am J Transl Res. 2014;6(2):114–8. PubMed PMID: 24489990; PubMed Central PMCID: PMC3902221.

11. Han JJ. FDA Modernization Act 2.0 allows for alternatives to animal testing. Artif Organs. 2023 Mar;47(3):449–50. doi:10.1111/aor.14503 PubMed PMID: 36762462.

12. Commissioner O of the. FDA [Internet]. FDA; 2025 [cited 2025 Apr 22]. FDA Announces Plan to Phase Out Animal Testing Requirement for Monoclonal Antibodies and Other Drugs. Available from: https://www.fda.gov/news-events/press-announcements/fda-announces-plan-phase-out-animal-testing-requirement-monoclonal-antibodies-and-other-drugs

13. Kondo J, Inoue M. Application of Cancer Organoid Model for Drug Screening and Personalized Therapy. Cells. 2019 May;8(5):5. doi:10.3390/cells8050470

14. Rep. Carter EL ‘Buddy’ [R G 1. Text - H.R.7248 - 118th Congress (2023-2024): FDA Modernization Act 3.0 [legislation] [Internet]. 2024 [cited 2024 Mar 25]. Located at: 2024-02-06. Available from: https://www.congress.gov/bill/118th-congress/house-bill/7248/text

15. Sato T, Stange DE, Ferrante M, Vries RGJ, Van Es JH, Van den Brink S, et al. Long-term expansion of epithelial organoids from human colon, adenoma, adenocarcinoma, and Barrett’s epithelium. Gastroenterology. 2011 Nov;141(5):1762–72. doi:10.1053/j.gastro.2011.07.050 PubMed PMID: 21889923.

16. van de Wetering M, Francies HE, Francis JM, Bounova G, Iorio F, Pronk A, et al. Prospective derivation of a living organoid biobank of colorectal cancer patients. Cell. 2015 May 7;161(4):933–45. doi:10.1016/j.cell.2015.03.053 PubMed PMID: 25957691; PubMed Central PMCID: PMC6428276.

17. Sato T, Vries RG, Snippert HJ, van de Wetering M, Barker N, Stange DE, et al. Single Lgr5 stem cells build crypt-villus structures in vitro without a mesenchymal niche. Nature. 2009 May;459(7244):262–5. doi:10.1038/nature07935

18. Vlachogiannis G, Hedayat S, Vatsiou A, Jamin Y, Fernández-Mateos J, Khan K, et al. Patient-derived organoids model treatment response of metastatic gastrointestinal cancers. Science. 2018 Feb 23;359(6378):920–6. doi:10.1126/science.aao2774 PubMed PMID: 29472484; PubMed Central PMCID: PMC6112415.

19. Esposito A, Agostini A, Quero G, Piro G, Priori L, Caggiano A, et al. Colorectal cancer patients-derived immunity-organoid platform unveils cancer-specific tissue markers associated with immunotherapy resistance. Cell Death Dis. 2024 Dec 4;15(12):878. doi:10.1038/s41419-024-07266-5 PubMed PMID: 39632825; PubMed Central PMCID: PMC11618451.

20. Zhao Y, Zhang B, Ma Y, Zhao F, Chen J, Wang B, et al. Colorectal Cancer Patient-Derived 2D and 3D Models Efficiently Recapitulate Inter- and Intratumoral Heterogeneity. Advanced Science. 2022;9(22):2201539. doi:10.1002/advs.202201539

21. Sakshaug BC, Folkesson E, Haukaas TH, Visnes T, Flobak Å. Systematic review: predictive value of organoids in colorectal cancer. Sci Rep. 2023 Oct 23;13(1):18124. doi:10.1038/s41598-023-45297-8 PubMed PMID: 37872318; PubMed Central PMCID: PMC10593775.

22. Tang Y, Wang T, Hu Y, Ji H, Yan B, Hu X, et al. Cutoff value of IC50 for drug sensitivity in patient-derived tumor organoids in colorectal cancer. iScience. 2023 Jul 21;26(7):107116. doi:10.1016/j.isci.2023.107116 PubMed PMID: 37426352; PubMed Central PMCID: PMC10329174.

23. Smabers LP, Wensink E, Verissimo CS, Koedoot E, Pitsa KC, Huismans MA, et al. Organoids as a biomarker for personalized treatment in metastatic colorectal cancer: drug screen optimization and correlation with patient response. J Exp Clin Cancer Res. 2024 Feb 27;43(1):61. doi:10.1186/s13046-024-02980-6

24. Piro G, Agostini A, Larghi A, Quero G, Carbone C, Esposito A, et al. Pancreatic Cancer Patient-Derived Organoid Platforms: A Clinical Tool to Study Cell- and Non-Cell-Autonomous Mechanisms of Treatment Response. Front Med (Lausanne). 2021 Dec 23;8:793144. doi:10.3389/fmed.2021.793144 PubMed PMID: 35004765; PubMed Central PMCID: PMC8733292.

25. Piro G, Carbone C, Agostini A, Esposito A, De Pizzol M, Novelli R, et al. CXCR1/2 dual-inhibitor ladarixin reduces tumour burden and promotes immunotherapy response in pancreatic cancer. Br J Cancer. 2023 Jan 19;128(2):331–41. doi:10.1038/s41416-022-02028-6 PubMed PMID: 36385556; PubMed Central PMCID: PMC9902528.

26. Raghavan S, Winter PS, Navia AW, Williams HL, DenAdel A, Lowder KE, et al. Microenvironment drives cell state, plasticity, and drug response in pancreatic cancer. Cell. 2021 Dec 9;184(25):6119–6137.e26. doi:10.1016/j.cell.2021.11.017 PubMed PMID: 34890551; PubMed Central PMCID: PMC8822455.

27. Ding S, Hsu C, Wang Z, Natesh NR, Millen R, Negrete M, et al. Patient-derived micro-organospheres enable clinical precision oncology. Cell Stem Cell. 2022 Jun 2;29(6):905–917.e6. doi:10.1016/j.stem.2022.04.006

28. Ooft SN, Weeber F, Dijkstra KK, McLean CM, Kaing S, van Werkhoven E, et al. Patient-derived organoids can predict response to chemotherapy in metastatic colorectal cancer patients. Sci Transl Med. 2019 Oct 9;11(513):eaay2574. doi:10.1126/scitranslmed.aay2574 PubMed PMID: 31597751.

29. Dekkers JF, Alieva M, Cleven A, Keramati F, Wezenaar AKL, van Vliet EJ, et al. Uncovering the mode of action of engineered T cells in patient cancer organoids. Nat Biotechnol. 2023 Jan;41(1):60–9. doi:10.1038/s41587-022-01397-w PubMed PMID: 35879361; PubMed Central PMCID: PMC9849137.

30. Harter MF, Recaldin T, Gerard R, Avignon B, Bollen Y, Esposito C, et al. Analysis of off-tumour toxicities of T-cell-engaging bispecific antibodies via donor-matched intestinal organoids and tumouroids. Nat Biomed Eng. 2024 Apr;8(4):345–60. doi:10.1038/s41551-023-01156-5 PubMed PMID: 38114742; PubMed Central PMCID: PMC11087266.

31. Dijkstra KK, Cattaneo CM, Weeber F, Chalabi M, Van De Haar J, Fanchi LF, et al. Generation of Tumor-Reactive T Cells by Co-culture of Peripheral Blood Lymphocytes and Tumor Organoids. Cell. 2018 Sep;174(6):1586–1598.e12. doi:10.1016/j.cell.2018.07.009

32. Cattaneo CM, Dijkstra KK, Fanchi LF, Kelderman S, Kaing S, Van Rooij N, et al. Tumor organoid–T-cell coculture systems. Nat Protoc. 2020 Jan;15(1):15–39. doi:10.1038/s41596-019-0232-9

33. Schnalzger TE, de Groot MH, Zhang C, Mosa MH, Michels BE, Röder J, et al. 3D model for CAR-mediated cytotoxicity using patient-derived colorectal cancer organoids. EMBO J. 2019 Jun 17;38(12):e100928. doi:10.15252/embj.2018100928 PubMed PMID: 31036555; PubMed Central PMCID: PMC6576164.

34. Price S, Bhosle S, Gonçalves E, Li X, McClurg DP, Barthorpe S, et al. A suspension technique for efficient large-scale cancer organoid culturing and perturbation screens. Sci Rep. 2022 Apr 2;12(1):5571. doi:10.1038/s41598-022-09508-y

35. Meric-Bernstam F, Hurwitz H, Raghav KPS, McWilliams RR, Fakih M, VanderWalde A, et al. Pertuzumab plus trastuzumab for HER2-amplified metastatic colorectal cancer (MyPathway): an updated report from a multicentre, open-label, phase 2a, multiple basket study. The Lancet Oncology. 2019 Apr 1;20(4):518–30. doi:10.1016/S1470-2045(18)30904-5 PubMed PMID: 30857956.

36. Janani B, Vijayakumar M, Priya K, Kim JH, Prabakaran DS, Shahid M, et al. EGFR-Based Targeted Therapy for Colorectal Cancer—Promises and Challenges. Vaccines (Basel). 2022 Mar 24;10(4):499. doi:10.3390/vaccines10040499 PubMed PMID: 35455247; PubMed Central PMCID: PMC9030067.

37. Li H, Durbin R. Fast and accurate long-read alignment with Burrows-Wheeler transform. Bioinformatics. 2010 Mar 1;26(5):589–95. doi:10.1093/bioinformatics/btp698 PubMed PMID: 20080505; PubMed Central PMCID: PMC2828108.

38. Van der Auwera GA, Carneiro MO, Hartl C, Poplin R, Del Angel G, Levy-Moonshine A, et al. From FastQ data to high confidence variant calls: the Genome Analysis Toolkit best practices pipeline. Curr Protoc Bioinformatics. 2013;43(1110):11.10.1-11.10.33. doi:10.1002/0471250953.bi1110s43 PubMed PMID: 25431634; PubMed Central PMCID: PMC4243306.

39. Chen X, Schulz-Trieglaff O, Shaw R, Barnes B, Schlesinger F, Källberg M, et al. Manta: rapid detection of structural variants and indels for germline and cancer sequencing applications. Bioinformatics. 2016 Apr 15;32(8):1220–2. doi:10.1093/bioinformatics/btv710 PubMed PMID: 26647377.

40. Roller E, Ivakhno S, Lee S, Royce T, Tanner S. Canvas: versatile and scalable detection of copy number variants. Bioinformatics. 2016 Aug 1;32(15):2375–7. doi:10.1093/bioinformatics/btw163 PubMed PMID: 27153601.

41. McLaren W, Gil L, Hunt SE, Riat HS, Ritchie GRS, Thormann A, et al. The Ensembl Variant Effect Predictor. Genome Biology. 2016 Jun 6;17(1):122. doi:10.1186/s13059-016-0974-4

42. Ewels PA, Peltzer A, Fillinger S, Patel H, Alneberg J, Wilm A, et al. The nf-core framework for community-curated bioinformatics pipelines. Nat Biotechnol. 2020 Mar;38(3):276–8. doi:10.1038/s41587-020-0439-x

43. Chen S, Zhang J, Shen M, Han X, Li S, Hu C, et al. p38 inhibition enhances TCR-T cell function and antagonizes the immunosuppressive activity of TGF-β. International Immunopharmacology. 2021 Sep 1;98:107848. doi:10.1016/j.intimp.2021.107848

44. Tharaux PL, Bukoski RC, Rocha PN, Crowley SD, Ruiz P, Nataraj C, et al. Rho Kinase Promotes Alloimmune Responses by Regulating the Proliferation and Structure of T Cells1. The Journal of Immunology. 2003 Jul 1;171(1):96–105. doi:10.4049/jimmunol.171.1.96

45. Nam GH, Lee EJ, Kim YK, Hong Y, Choi Y, Ryu MJ, et al. Combined Rho-kinase inhibition and immunogenic cell death triggers and propagates immunity against cancer. Nat Commun. 2018 Jun 4;9(1):2165. doi:10.1038/s41467-018-04607-9

46. Fujii M, Matano M, Toshimitsu K, Takano A, Mikami Y, Nishikori S, et al. Human Intestinal Organoids Maintain Self-Renewal Capacity and Cellular Diversity in Niche-Inspired Culture Condition. Cell Stem Cell. 2018 Dec 6;23(6):787–793.e6. doi:10.1016/j.stem.2018.11.016 PubMed PMID: 30526881.

47. Yui S, Azzolin L, Maimets M, Pedersen MT, Fordham RP, Hansen SL, et al. YAP/TAZ-Dependent Reprogramming of Colonic Epithelium Links ECM Remodeling to Tissue Regeneration. Cell Stem Cell. 2018 Jan 4;22(1):35–49.e7. doi:10.1016/j.stem.2017.11.001 PubMed PMID: 29249464; PubMed Central PMCID: PMC5766831.

48. Serra D, Mayr U, Boni A, Lukonin I, Rempfler M, Challet Meylan L, et al. Self-organization and symmetry breaking in intestinal organoid development. Nature. 2019 May;569(7754):66–72. doi:10.1038/s41586-019-1146-y PubMed PMID: 31019299; PubMed Central PMCID: PMC6544541.

49. Pinna CMA, Rakhimova K, Zhang J, Bills L, Tee L, Blanc C, et al. A step-by-step low-matrix suspension co-culture method to screen T cell engager activity in patient-derived… [Internet]. 2026 Sep 21 [cited 2026 Sep 21]. Available from: https://www.protocols.io/view/a-step-by-step-low-matrix-suspension-co-culture-me-kxygxjd1dl8j/v1

