## Supplementary Document S1 for "A low-matrix organoid-T cell co-culture platform for functional evaluation of T cell engagers in patient-derived colorectal cancer organoids"

**SUPPLEMENTARY TABLES**

|  | **OM (Organoids in**  **house Media)** | **RPMI & 10% FBS (v/v)** | **Immunocult** | **RPMI 10% FBS: OM**  **(80:20 v/v)** | **RPMI 10% FBS: OM -SB**  **(80:20 v/v)** | **OM -SB/-Y27632: Immunocult (50:50 v/v)** | **Intesticult:**  **Immunocult**  **(50:50 v/v)** |
| --- | --- | --- | --- | --- | --- | --- | --- |
| Intesticult (IntestiCult Organoid Growth Medium (Human) | - | - | - | - | - | - | 1X |
| Immunocult (ImmunoCult-XF T Cell Expansion Medium) | - | - | 1X | - | - | 1X | 1X |
| Advanced DMEM/F12 | 1X | - | - | 1X | 1X | 1X | - |
| RPMI 1640 Medium | - | 1 X | - | 1 X | 1 X | 1 X | - |
| Fetal Bovine Serum (FBS) | - | 10% (v/v) | - | 10% (v/v) | 10% (v/v) | 10% (v/v) | - |
| GlutaMax | 1 X | - | - | 1 X | 1 X | 1 X | - |
| Hepes | 10 mM | - | - | 10 mM | 10 mM | 10 mM | - |
| N-2 MAX Supplement | 1 X | - | - | 1 X | 1 X | 1 X | - |
| Human R-Spondin1 | 500 ng/mL | - | - | 500 ng/mL | 500 ng/mL | 500 ng/mL | - |
| Human Noggin | 100 ng/mL | - | - | 100 ng/mL | 100 ng/mL | 100 ng/mL | - |
| B27 supplement | 1 X | - | - | 1 X | 1 X | 1 X | - |
| N-Acetyl-Cysteine | 1.25 mM | - | - | 1.25 mM | 1.25 mM | 1.25 mM | - |
| Nicotinamide | 10 mM | - | - | 10 mM | 10 mM | 10 mM | - |
| Recombinant Human EGF | 50 ng/mL | - | - | 50 ng/mL | 50 ng/mL | 50 ng/mL | - |
| A83-01 | 500 nM | - | - | 500 nM | 500 nM | 500 nM | - |
| Prostaglandin E2 | 10 nM | - | - | 10 nM | 10 nM | 10 nM | - |
| Gastrin 1 human | 10 nM | - | - | 10 nM | 10 nM | 10 nM | - |
| SB 202190 | 30 mM | - | - | 30 mM | - | - | - |
| Rho kinase inhibitor Y27632^a^ | 10 µM | - | - | 10 µM | 10 µM | - | 10 µM |

**Table S1. Overview of all screened media variants, including individual components and their final concentrations (Related to Figure 1). ^a^Y‑27632 is added to the culture medium only during the recovery phase after single‑cell organoid seeding and is kept for 24–48 hours following passaging.**

| **Donor** | **HLA- ABC class I** | | | | | |
| --- | --- | --- | --- | --- | --- | --- |
|  | **HLA_A1** | **HLA_A2** | **HLA_B1** | **HLA_B2** | **HLA_C1** | **HLA_C2** |
| 804C | A*01:01 | A*03:01 | B*18:01 | B*57:01 | C*07:01 | C*06:02 |
| 702C | A*26:01 | A*26:01 | B*38:01 | B*55:01 | C*03:03 | C*12:03 |
| 105C | A*02:01 | A*02:02 | B*45:01 | B*58:02 | C*06:02 | C*16:01 |
| 054 | A*23:01 | A*32:01 | B*15:03 | B*44:02 | C*02:10 | C*05:01 |
| LMX129 | A*02:01:01 | A*03:01:01 | B*07:02:01 | B*51:01:01 | C*01:02:01 | C*07:02:01 |
| **Donor** | **HLA_DQA1_1** | **HLA_DQA1_2** | **HLA_DPB1_1** | **HLA_DPB1_2** | **HLA_DQB1_1** | **HLA_DQB1_2** |
| 804C | - | - | DPB1*04:01 | DPB1*04:02 | DQB1*03:03 | DQB1*06:02 |
| 702C | DQA1*02:01 | DQA1*05:01 | DPB1*04:01 | DPB1*04:01 | DQB1*02:01 | DQB1*03:01 |
| 105C | DQA1*02:01 | DQA1*05:01 | DPB1*04:02 | DPB1*85:01 | DQB1*02:01 | DQB1*03:01 |
| 054 | - | - | DPB1*02:01 | DPB1*04:02 | DQB1*02:01 | DQB1*06:02 |
| LMX129 | DQA1*01:03:01 | DQA1*02:01:01 | DPB1*02:01:02 | DPB1*04:02:01 | DQB1*03:03:02 | DQB1*06:03:01 |
| **Donor** | **HLA-DRB** | | | |  |  |
| 804C | DRB1*07:01 | DRB1*15:01 | DRB4*01:01 | DRB5*01:01 |  |  |
| 702C | DRB1*07:01 | DRB1*12:01 | DRB3*02:02 | DRB4*01:01 |  |  |
| 105C | DRB1*07:01 | DRB1*11:02 | DRB3*02:02 | DRB4*01:01 |  |  |
| 054 | DRB1*07:01 | DRB1*11:01 | DRB3*02:02 | DRB4*01:01 |  |  |
| LMX129 | DRB1*07:01:01 | DRB1*13:01:01 | DRB3*02:02:01 | DRB4*01:03:01N |  |  |

**Table S2. Human healthy leukopaks donors’ HLA Locus, Chromosome and Allele (Related to Figure 3).**

| **SAMPLE ID** | **S403376** | **S302389** | **S302411** | **S366557** | **S315571** | **S345653** | **COLO**  **151** | **COLO 143** | **COLO 240** | **COLO**  **250** | **COLO 298** | **COLO 302** | **COLO**  **338** | **COLO 356** | **COLO**  **360** | **COLO**  **376** | **COLO 378** | **COLO 388** | **COLO**  **394** | **COLO 434** | **COLO**  **454** |
| --- | --- | --- | --- | --- | --- | --- | --- | --- | --- | --- | --- | --- | --- | --- | --- | --- | --- | --- | --- | --- | --- |
| Age | 51 | 81 | 74 | 75 | 61 | 55 | 78 | 70 | 64 | 69 | 60 | 76 | 83 | 84 | 64 | 74 | 72 | 57 | 65 | 74 | 64 |
| Gender | M | M | F | F | M | F | M | F | F | F | F | F | M | F | F | M | M | M | M | M | M |
| Anatomic Site | Rectum | Right | Right | Right | Rectum | Rectum | Right | Right | Rectum | Rectum | Right | Right | Right | Left | Left | Right | Left | Rectum | Rectum | Left | Left |
| Tstage | 3 | 4 | 3 | 3 | 1 | 2 | 3 | 3 | 3 | 4 | 4 | 3 | 3 | 4 | 3 | 3 | 3 | 4 | 4 | 3 | 4 |
| Nstage | 0 | 2 | 2 | 0 | 0 | 0 | 0 | 1 | 0 | 0 | 0 | 0 | 1 | 1 | 0 | 0 | 0 | 0 | 0 | 2 | 1 |
| Mstage | 0 | 0 | 1 | 0 | 0 | 0 | 0 | 0 | 0 | 0 | 0 | 0 | 0 | 0 | 0 | 0 | 0 | 1 | 0 | 0 | 0 |
| Differentiation | Poor | Mod | Mod | Poor | Mod | Mod | Mod | Mod | Mod | Mod | Mod | Poor | Mod | Mod | Mod | Mod | Mod | Mod | Mod | Mod | Mod |
| EMVI | 1 | 1 | 1 | 0 | 0 | 1 | 0 | 1 | 0 | 0 | 1 | 0 | 1 | 1 | 1 | 0 | 1 | 0 | 1 | 1 | 1 |
| LVI | 0 | 0 | 0 | 0 | 0 | 0 | 0 | 0 | 1 | 0 | 1 | 0 | 1 | 1 | 1 | 0 | 0 | 0 | 0 | 1 | 1 |
| PNI | 0 | 0 | 0 | 1 | 0 | 0 | 0 | 0 | 1 | 0 | 1 | 0 | 1 | 0 | 0 | 0 | 0 | 0 | 1 | 0 | 1 |
| CMS Classifier | CMS1 | CMS2 | CMS3 | CMS1 | CMS1 | CMS4 | CMS4 | CMS4 | CMS3 | CMS3 | CMS2 | CMS1 | CMS2 | CMS3 | CMS4 | CMS1 | CMS4 | CMS3 | CMS4 | CMS2 | CMS2 |
| PreOp Treatment | LCCRT | None | None | None | None | LCCRT | None | None | RAPIDO | SCRT | None | None | None | None | None | None | FOLFOX | None | RAPIDO | FOLFOX | None |
| **Mutations** |  |  |  |  |  |  |  |  |  |  |  |  |  |  |  |  |  |  |  |  |  |
| KRAS | G13R |  |  | G13R |  | G12D |  |  |  |  | G12D |  |  | G12D |  | G13D |  | G12D |  | G13D |  |
| APC | Y1376* V1822D | Q1338Afs*4 V1822D | Q1338Afs*4 Q1378* | V1822D A921S | K1007* N1797Ifs3* | E1322Sfs*6 V1479Sfs*28 V1822D | Q1152* T1556Nfs*3 | Y935* R1450* |  | T1556Nfs*3 T909Nfs*3 Y1031H | S1356* | F801Lfs*19 | R1114* |  | P1243Gfs*10 P1993A | X470_splice D917* | R216* S1346* | Q1378* R554* | T1301Nfs*14 | S1315* | R805* |
| TP53 | R175H G245D | G245D G245S | R175H R248Q | G245D |  | P191del |  | R248Q |  |  |  |  | X332_splice | R248W | E204* |  | E180* | E271K | R248W | R110L | L93Vfs*55 |
| BRAF |  |  | V600E |  |  |  | V600E |  |  | D594G |  | V600E |  |  |  |  |  |  |  |  |  |
| CTNNB1 |  | L159R R474Q |  |  |  | F683Qfs*9 |  |  | S45F |  |  |  |  | S45F |  |  |  |  |  |  |  |
| POLD1 |  |  |  |  |  |  |  |  | L58R |  |  |  |  |  |  |  | D496A |  |  |  |  |
| POLE |  |  |  |  |  |  |  |  |  |  |  |  |  |  |  |  |  |  |  |  | A252V |
| **RNA Expression (in TPM)** |  |  |  |  |  |  |  |  |  |  |  |  |  |  |  |  |  |  |  |  |  |
| EGFR | 42.9 | 12.94 | 9.12 | 38.09 | 4.39 | 35.99 | 23.07 | 8.57 | 5.66 | 14.82 | 1.32 | 10.02 | 27.05 | 4.15 | 13.61 | 39.8 | 19.1 | 6.89 | 11.96 | 9.35 | 16.65 |
| ERBB2 | 84.97 | 39.11 | 39.03 | 76.42 | 18.09 | 82.8 | 177.39 | 67.64 | 41.41 | 170.21 | 32.1 | 48.68 | 281.67 | 33.71 | 49.56 | 264.82 | 66.87 | 82.12 | 93.48 | 142 | 171.28 |
| MSI status | MSS | MSS | MSS | MSI | MSI | MSS | MSS | MSS | MSI | MSS | MSS | MSI | MSS | MSS | MSS | MSI | MSS | MSS | MSS | MSS | MSS |
| TMB | 5.45 | 8.24 | 3.55 | 111.6 | 65.4 | 9.84 | 13.95 | 11.41 | 121.72 | 9.01 | 4.56 | 123.54 | 6.75 | 10.82 | 4.84 | 115.11 | 5.24 | 10.73 | 11.45 | 6.07 | 8.41 |

**Table S3. Patients clinical references and mutations of the specimens from which the CRC organoids models were derived (Related to Figure 4).** M, male; F, female; Differentiation: P, Poor; Mod, Moderate; MSS, Microsatellite Stable; MSI, Microsatellite Instable; TMB, Tumour mutational burden.

| **ORGANOID** | **HLA- ABC class I** | | | | | |
| --- | --- | --- | --- | --- | --- | --- |
|  | **HLA_A1** | **HLA_A2** | **HLA_B1** | **HLA_B2** | **HLA_C1** | **HLA_C2** |
| **S366557** | A*01:01:01:01 | A*01:01:01:01 | B*37:01:01:09 | B*08:01:01:01 | C*06:02:01:01 | C*07:01:01:12 |
| **COLO151** | A*24:02:102:01 | A*24:02:10 | B*07:02:01:01 | B*07:02:01:01 | C*07:02:01:03 | C*07:02:01:03 |
| **COLO250** | A*26:01:01G | A*33:03:01G | B*08:01:01G | B*58:01:01G | C*03:02:01G | C*07:02:01G |
| **COLO298** | A*01:01:01G | A*24:02:01G | B*35:01:01G | B*58:01:01G | C*03:16 | C*04:01:01G |
| **S345653** | A*24:02:01G | A*24:149 | B*35:03:01G | B*35:03:07 | C*12:03:01G | C*12:03:01G |
| **S315571** | A*01:01:01:01 | A*01:01:01:01 | B*08:01:01:01 | B*37:01:01:01 | C*06:04:01 | C*06:02:01:01 |
| **ORGANOID** | **HLA_DPA1_1** | **HLA_DPA1_2** | **HLA_DPB1_1** | **HLA_DPB1_2** | **HLA_DQA1_1** | **HLA_DQA1_2** |
| **S366557** | DPA1*01:03:01:04 | DPA1*01:03:01:03 | DPB1*25:01 | DPB1*04:01:01:15 | DQA1*05:05:01:01 | DQA1*01:02:01:01 |
| **COLO151** | DPA1*02:01:02:02 | DPA1*02:01:02:02 | DPB1*01:01:01:01 | DPB1*01:01:01:01 | DQA1*01:03:01:02 | DQA1*01:03:01:02 |
| **COLO250** | DPA1*01:03:01G | DPA1*01:03:01G | DPB1*02:01:02G | DPB1*04:01:01G | DQA1*01:02:01G | DQA1*01:03:01G |
| **COLO298** | DPA1*01:03:01G | DPA1*01:03:01G | DPB1*02:01:02G | DPB1*04:01:01G | DQA1*01:01:01G | DQA1*01:03:01G |
| **S345653** | DPA1*02:02:05 | DPA1*02:02:02 | DPB1*26:01:02 | DPB1*26:01:02 | - | - |
| **S315571** | DPA1*01:03:01:04 | DPA1*01:03:01:04 | DPB1*16:01:01:02 | DPB1*04:01:01:05 | DQA1*05:05:01:19 | DQA1*05:05:01:01 |
| **ORGANOID** | **HLA_DRB1_1** | **HLA_DRB1_2** | **HLA_DRB3_1** | **HLA_DRB3_2** | **HLA_DRB4_1** | **HLA_DRB4_2** |
| **S366557** | DRB1*11:03:01 | DRB1*15:01:01:05 | - | - | - | - |
| **COLO151** | DRB1*13:01:01:01 | DRB1*13:02:01:02 | - | - | - | - |
| **COLO250** | DRB1*13:01:01G | DRB1*13:02:01 | DRB3*01:01:02G | DRB3*03:01:01G | DRB4*03:01N | DRB4*03:01N |
| **COLO298** | DRB1*13:01:01G | DRB1*01:71 | DRB3*03:01:01G | DRB3*03:01:01G | DRB4*03:01N | DRB4*03:01N |
| **S345653** | DRB1*11:01:01G | DRB1*11:23:02 | DRB3*02:02:01G | DRB3*02:02:01G | DRB4*03:01N | DRB4*03:01N |
| **S315571** | DRB1*11:01:01:01 | DRB1*11:01:02 | - | - | - | - |
| **ORGANOID** | **HLA_DQA1_1** | **HLA_DQA1_2** | **HLA_DQB1_1** | **HLA_DQB1_2** |  |  |
| **S366557** | - | - | - | DQB1*03:01:01:18 |  |  |
| **COLO151** | - | - | DQB1*06:03:23 | - |  |  |
| **COLO250** | DQA1*01:02:01G | DQA1*01:03:01G | DQB1*06:03:01G | DQB1*06:18:02 |  |  |
| **COLO298** | DQA1*01:01:01G | DQA1*01:03:01G | DQB1*06:03:01G | DQB1*05:01:14 |  |  |
| **S345653** | DQA1*05:01:01G | DQA1*05:01:01G | DQB1*03:01:01G | DQB1*03:01:01G |  |  |
| **S315571** | - | - | DQB1*03:01:01:03 | - |  |  |

**Table S4. CRC Organoids Locus, Chromosome and Allele HLA typing (Related to Figure 4).**

| **REAGENT OR RESOURCE** | **SOURCE** | **IDENTIFIER** |
| --- | --- | --- |
| **Antibodies** | | |
| CD4, Mouse anti-Human | BioLegend | Cat#344624 |
| CD8a, Mouse anti-Human | BioLegend | Cat#344712 |
| CD45RA, Mouse anti-Human | BioLegend | Cat#304140 |
| CD45RO, Mouse anti-Human | BioLegend | Cat#304206 |
| CCR7, Mouse anti-Human | BioLegend | Cat#353206 |
| CD25, Mouse anti-Human | BioLegend | Cat#302634 |
| PD-1, Mouse anti-Human | BioLegend | Cat#329916 |
| CD137, Mouse anti-Human | BioLegend | Cat#309826 |
| CD69, Mouse anti-Human | BioLegend | Cat#310910 |
| TIM-3, Mouse anti-Human | BioLegend | Cat#345030 |
| CD45, Mouse anti-Human | BioLegend | Cat#368540 |
| CD326, Mouse anti-Human | BioLegend | Cat#324206 |
| PE anti-human EGFR | BioLegend | Cat#352904 |
| PE anti-human CD340 (erbB2/HER-2) | BioLegend | Cat#324405 |
| Isotype Control PE Mouse IgG1, κ Isotype Ctrl (FC) | BioLegend | Cat#400114 |
| CoraLite Plus 488-conjugated HER2/ErbB2 | Proteintech | Cat#CL488-60311 |
| CoraLite Plus 647-conjugated Phospho-EGFR (Tyr1197) | Proteintech | Cat#CL647-84906 |
| eBioscience Fixable Viability Dye eFluor 780 | Invitrogen | Cat#65-0865-14 |
| Zombie Green Fixable Viability Kit | BioLegend | Cat#423112 |
| EGFR-TCE | AstraZeneca | N/A |
| HER2-TCE | AstraZeneca | N/A |
| Isotype-TCE | AstraZeneca | N/A |
| Bovine Albumin Fraction V (7.5 % solution) | ThermoFisher Scientific | Cat#15260037 |
| Triton X-100 | ThermoFisher Scientific | Cat# A16046.AP |

| **Biological samples** |
| --- |

| Healthy donor leukopak 211282804C (804C) | Stemcell Technologies | Cat#200-0130 |
| --- | --- | --- |
| Healthy donor leukopak 220980702C (702C) | Stemcell Technologies | Cat#200-0130 |
| Healthy donor leukopak 220980105C (105C) | Stemcell Technologies | Cat#200-0130 |
| Healthy donor leukopak 2212409054 (054) | Stemcell Technologies | Cat#200-0130 |
| Healthy donor leukopak LMX100-0001129 (LMX_129) | BioIVT | Cat#1711131 |
| **Chemicals, peptides, and recombinant proteins** | | |
| Ultra Comp eBeads | ThermoFisher Scientific | Cat#01-2222-42 |
| EDTA | Invitrogen | Cat#10135423 |
| ImmunoCult-XF T Cell Expansion Medium | Stemcell Technologies | Cat#10981 |
| DNase I Solution (1 mg/mL) | Stemcell Technologies | Cat#7900 |
| Human Recombinant IL-2 (CHO-expressed) | Stemcell Technologies | Cat#78036.3 |
| ImmunoCult Human CD3/CD28 T Cell Activator | Stemcell Technologies | Cat#10971 |
| EasySep Buffer | Stemcell Technologies | Cat#20144 |
| NucBlue Fixed Cell ReadyProbes Reagent (DAPI) | ThermoFisher Scientific | Cat# R37606 |
| CellTracker Deep Red | Invitrogen | Cat# C34565 |
| Incucyte Caspase-3/7 Green Dye for Apoptosis | Sartorius | Cat#4440 |
| Advanced DMEM/F12 | Fisher | Cat#12634028 |
| DPBS | Gibco | Cat#14190144 |
| Fetal Bovine Serum (FBS) | Gibco | Cat#17593595 |
| MACS Tissue storage solution | Miltenyi | Cat#130-100-008 |
| Anti-Adherence Rinsing Solution (Stemcell Technologies | Stemcell Technologies | Cat#07010 |
| IntestiCult Organoid Growth Medium (Human) | Stemcell Technologies | Cat#06010 |
| GlutaMax | Invitrogen | Cat#35050038 |
| Hepes | Invitrogen | Cat#15630-056 |
| N-2 MAX Supplement | Bio-techne | Cat#AR009 |
| B27 supplement | Gibco | Cat#17504-44 |
| Human R-Spondin 1 Recombinant Protein, PeproTech | Gibco | Cat#120-38 |
| Human Noggin Recombinant Protein, PeproTech | Gibco | Cat#120-10C |
| N-Acetylcysteine | Sigma | Cat# A9165 |
| Nicotinamide | Sigma | Cat# N0636 |
| Human EGF, Animal-Free Recombinant Protein, PeproTech | Gibco | Cat# AF-100-15 |
| A83-01 | Tocris | Cat#2939-10 |
| SB 202190 | Sigma | Cat# S7067 |
| Prostaglandin E2 (PGE2) | Tocris | Cat#2296 |
| Gastrin 1 human | Bio-techne | Cat#3006/1 |
| Rho kinase inhibitor Y-27632 dihydrochloride | R and D systems | Cat#1254 |
| Cultrex UltiMatrix RGF Basement Membrane Extract (UltiMatrix BME) | Bio-techne | Cat#BME001-10 |
| Primocin | Invitrogen | Cat#ant-pm-1 |
| MycoStrip - Mycoplasma Detection Kit | Invitrogen | Cat#rep-mys-50 |
| TrypLE express | Invitrogen | Cat#12605028 |
| Collagenase II | Gibco | Cat#17101015 |
| Dispase II | Gibco | Cat#17105041 |
| **Critical commercial assays** | | |
| Quantibrite PE Phycoerythrin Fluorescence Quantitation Kit (RUO (GMP)) | BD | Cat#340495 |
| Human EasySep T Cell Iso Kit | Stemcell Technologies | Cat#100-0695 |
| Ligation sequencing kit v14 | Oxford Nanopore Technology | Cat# SQK-LSK114 |
| Direct RNA Sequencing Kit | Oxford Nanopore Technologies | Cat#SQK-RNA002 |
| PromethION Flow Cells Packs (DNA) | Oxford Nanopore Technologies | Cat#FLO-PRO114M |
| PromethION Flow Cells Packs (RNA) | Oxford Nanopore Technologies | Cat#FLO-PRO004RA |
| LEGENDplex Human CD8/NK Panel (13-plex) w/ VbP V02 | BioLegend | Cat#741187 |
| Monarch Spin RNA Isolation Kit (Mini) | NEB | Cat#T2110 |
| Monarch Genomic DNA Purification Kit | NEB | Cat#T3010 |
| **Software and algorithms** | | |
| Prism10.4.1 | GraphPad | N/A |
| Gen5 software 3.16.10 | Agilent | N/A |
| NovoExpress 1.6.2 | Agilent | N/A |
| BioLegend LEGENDplex | BioLegend | https://legendplex.qognit.com |
| **Equipment** | | |
| Flow cytometer Novocyte Panteon | Agilent | N/A |
| Easy 250 EasySep magnet | Stemcell Technologies | Cat#100-0821 |
| EVOS XL Core Cell Imaging System | Thermo Fisher Scientific | Cat# AMEX1100 |
| BioTek Cytation 5 cell imaging multimode reader | Agilent | N/A |
| BioTek BioSpa 8 Automated Incubator | Agilent | N/A |
| TapeStation 4200 system | Agilent | N/A |
| Countess 3 Automated Cell Counter | Invitrogen | N/A |
| Countess Cell Counting Chamber Slides | Thermo Fisher Scientific | Cat#C10228 |
| Sonicator | Covaris | Cat#E220 |
| g-Tubes | Covaris | Cat#520079 |
| PromethION 24 A100 | Oxford Nanopore Technologies | N/A |

**Table S5. Key Resources Table (Related to Supplementary Methods).** Summary of all biological materials, antibodies, reagents, commercial assays, sequencing resources, software, and instrumentation used in this study, including source information and catalog identifiers to support reproducibility of the organoid-T cell co-culture workflow and immune engager screening platform.

**SUPPLEMENTARY METHODS**

A detailed, step-by-step protocol used in this study is available on Protocols.io**: *A step-by-step low-matrix suspension co-culture method to screen T cell engager activity in patient-derived organoids*** (DOI: <https://doi.org/10.17504/protocols.io.kxygxjd1dl8j/v1>) (49).

**Patient samples and ethics**

Patients undergoing surgical resection for colorectal cancer (CRC) at University Hospitals Birmingham NHS Foundation Trust, UK, were recruited under approved ethical protocols (North West-Haydock Research Ethics Committee; REC reference 20/NW/0001, previous REC reference 15/NW/0079; HBRC sub-approval 17-287). Written informed consent was obtained from all participants. Clinicopathological data were collected through the University of Birmingham Human Biomaterials Resource Centre.

**CRC organoid generation and culture**

After collection, colorectal cancer (CRC) specimens were stored at 4 °C in MACS Tissue Storage Solution (Miltenyi, Cat# 130-100-008) and processed within 24 hours for organoid derivation, following previously published CRC organoid culture protocols (15-17). Briefly, samples were mechanically fragmented and enzymatically digested in tumour digestion buffer consisting of 75 U/mL Collagenase II (Gibco, Cat#17101015), 125 µg/mL Dispase II (Gibco, Cat#17105041), 10 µM Y‑27632 (R&D Systems, Cat#1254), and 0.1 mg/mL Primocin (InvivoGen, Cat#ant‑pm‑1) in DMEM for 30–40 minutes at 37 °C.

Following digestion, epithelial clusters and single cells—visualised using the EVOS XL Core Cell Imaging System (Thermo Fisher Scientific, Cat#AMEX1100)—were washed in rinsing solution (DMEM containing 10% FBS [Gibco, Cat#17593595] and 0.1 mg/mL Primocin [InvivoGen, Cat#ant‑pm‑1]), filtered through a 70 µm cell strainer, pelleted, washed with ice‑cold DPBS (Gibco, Cat#14190144), and resuspended in UltiMatrix BME (Bio‑Techne, Cat#BME001‑10) at 50 µL per well.

Organoid domes were incubated at 37 °C with 5% CO₂ for 15–20 minutes to allow UltiMatrix polymerisation before addition of IntestiCult Human Organoid Growth Medium (STEMCELL Technologies, Cat#06010) supplemented with 10 µM Y‑27632 (R&D Systems, Cat#1254) and 0.1 mg/mL Primocin (InvivoGen, Cat#ant‑pm‑1). Y‑27632 was included only during the first week of culture, and Primocin during the first month.

Organoids were expanded under standard conditions, routinely tested for mycoplasma contamination using the MycoStrip Mycoplasma Detection Kit (InvivoGen, Cat#rep‑mys‑50), and cryopreserved at early passage numbers (< 10) for long‑term storage in liquid nitrogen.

A subset of CRC organoid models used in this study was generated by the Wellcome Trust Sanger Institute as part of the Human Cancer Models Initiative (HCMI), including COLO151, COLO143, COLO240, COLO250, COLO298, COLO302, COLO338, COLO356, COLO360, COLO376, COLO378, COLO388, COLO394, COLO434, and COLO454. Details of organoid models, matched clinical annotations, and HLA typing are provided in **Supplementary Tables 3 and 4**. Microsatellite stability status was confirmed by mismatch repair immunohistochemistry and whole‑genome sequencing analysis using MSISensor (52). All organoid models were authenticated by total RNA sequencing, whole‑genome sequencing, and Oxford Nanopore Technologies sequencing prior to experimental use.

CRC organoids were routinely passaged every 6–7 days. For passaging, organoids were harvested in ice‑cold DPBS (Gibco, Cat#14190144) and dissociated into single cells using pre‑warmed TrypLE Express (Invitrogen, Cat#12605028) supplemented with 10 µM Y‑27632 (R&D Systems, Cat#1254), combined with gentle mechanical trituration until predominantly single cells were observed. Enzymatic dissociation was terminated by dilution with ice‑cold DPBS, followed by centrifugation at 400 × g for 4 minutes at 4 °C.

Cell pellets were resuspended in UltiMatrix BME (Bio-Techne, Cat# BME001-10) and plated as domes in 24-well plates (80,000-100,000 cells per dome). Following polymerisation, domes were overlaid with IntestiCult Human Organoid Growth Medium (STEMCELL Technologies, Cat# 06010) supplemented with Y-27632 (R&D Systems, Cat# 1254) for the initial 2-3 days of culture.

For cryopreservation, organoids at early passage and in an undifferentiated growth state were gently dissociated, pelleted, and resuspended in freezing medium consisting of 80% Fetal Bovine Serum (Gibco, Cat#17593595) and 20% dimethyl sulfoxide (DMSO). Samples were transferred to labelled cryovials and frozen using a controlled‑rate freezing container at −80 °C before long‑term storage in liquid nitrogen. For recovery, cryovials were rapidly thawed in a 37 °C water bath, diluted in pre‑warmed DMEM to remove residual DMSO, and centrifuged. Organoids were re‑embedded in UltiMatrix BME (Bio‑Techne, Cat#BME001‑10), plated as domes in pre‑warmed 24‑well plates, and cultured in IntestiCult Human Organoid Growth Medium (STEMCELL Technologies, Cat#06010) as described above.

**T cells isolation and activation**

Peripheral blood mononuclear cells (PBMCs) were isolated from healthy human donor leukopaks obtained from STEMCELL Technologies (STEMCELL Technologies, Cat# 200-0130) or BioIVT (BioIVT, Cat# 1711131) according to the manufacturers’ instructions. Briefly, cryopreserved leukopaks were rapidly thawed in a 37 °C water bath and diluted 1:1 with wash buffer consisting of PBS supplemented with 10% (v/v) FBS (Gibco, Cat# 17593595). Cell suspensions were aliquoted into 50 mL conical tubes and centrifuged at 300 × g for 5 minutes at room temperature. Pellets were resuspended in DNase I solution (0.1 mg/mL; STEMCELL Technologies, Cat# 7900) and incubated for 15 minutes to reduce cell aggregation. Cells were then diluted with wash buffer, passed through a 40 µm cell strainer, and centrifuged at 300 × g for 10 minutes.

PBMCs were counted using a Countess 3 Automated Cell Counter (Thermo Fisher Scientific) and either cryopreserved in freezing medium consisting of 90% (v/v) FBS and 10% (v/v) DMSO or used immediately for CD3+ T cell isolation. CD3+ T cells were purified by immunomagnetic negative selection using the EasySep Human T Cell Isolation Kit (STEMCELL Technologies, Cat# 100-0695) and an EasySep Magnet (STEMCELL Technologies, Cat# 100-0821), according to the manufacturer's instructions.

**CRC organoids, CD3+ T cells, and proof‑of‑concept T cell engager (POC‑TCE) co‑cultures**

Cryopreserved CD3+ T cells were thawed and activated for 7 days in ImmunoCult‑XF T cell Expansion Medium (STEMCELL Technologies, Cat# 10981) supplemented with Human CD3/CD28 T cell Activator (25 µL/mL; STEMCELL Technologies, Cat# 10971). Following initial expansion, activated CD3+ T cells were harvested and cultured for an additional 3 days in fresh ImmunoCult‑XF medium supplemented with recombinant human interleukin‑2 (IL‑2) (10 ng/mL; STEMCELL Technologies, Cat# 78036.3). Prior to co‑culture, activated CD3+ T cells were washed to remove residual CD3/CD28 stimulation and cytokines.

CRC organoids were dissociated into single cells and seeded 5 days before co‑culture at a density of 10,000 cells per well in 96‑well flat‑bottom plates pre‑treated using Anti‑Adherence Rinsing Solution (STEMCELL Technologies, Cat# 07010). Organoids were cultured in an in-house CRC organoid medium (O.M.; lacking SB202190 and Y-27632) consisting of Advanced DMEM/F12 (Fisher, Cat# 12634028) supplemented with HEPES (10 mM; Invitrogen, Cat# 15630‑056), GlutaMAX™ (1×; Invitrogen, Cat# 35050038), N‑2 MAX Supplement (1×; Bio‑Techne, Cat# AR009), B27 Supplement (1×; Gibco, Cat# 17504‑44), N‑acetylcysteine (1.25 mM; Sigma, Cat# A9165), human recombinant R‑Spondin‑1 (500 ng/mL; Gibco, Cat# 120‑38), human recombinant Noggin (100 ng/mL; Gibco, Cat# 120‑10C), nicotinamide (10 mM; Sigma, Cat# N0636), human recombinant EGF (50 ng/mL; Gibco, Cat# AF‑100‑15), gastrin‑1 (10 nM; Bio‑Techne, Cat# 3006/1), A83‑01 (500 nM; Tocris, Cat# 2939‑10), and prostaglandin E₂ (PGE₂) (10 nM; Tocris, Cat# 2296), supplemented with

1% (v/v) UltiMatrix BME (Bio‑Techne, Cat# BME001‑10). Edge wells were filled with PBS to minimise evaporation. The culture medium was adapted from previously published CRC organoid culture protocols (15-17).

On the day of co‑culture, POC‑TCE molecules targeting EGFR or HER2, together with isotype and vehicle controls, were prepared in ImmunoCult‑XF medium at final concentrations ranging from 0.01 to 100 nM. CD3+ T cells were added to organoid cultures in a final volume of 200 µL per well, combining in-house CRC organoid medium (O.M.; lacking SB202190 and Y-27632) (100 µL) and ImmunoCult‑XF medium (100 µL) at a 1:1 ratio. Unstimulated CD3+ T cells were thawed and maintained without activation on the co-culture day. Effector‑to‑target ratios were set to 10:1 for unstimulated CD3+ T cells (100,000 T-cells to 10,000 organoid cells) and 2:1 for activated CD3+ T cells (20,000 T-cells to 10,000 organoid cells). Co‑cultures were incubated at 37 °C with 5% CO₂ without medium exchange until the indicated experimental endpoints.

**Flow cytometry and antibodies**

Prior to co‑culture experiments, isolated CD3+ T cells were phenotyped by flow cytometry. Cells were stained with Fixable Viability Dye eFluor 780 (Invitrogen, Cat#65‑0865‑14; 1 µL dye per 1 mL cell suspension) for 30 minutes at 4 °C in the dark, followed by surface staining for 30 minutes at 4 °C, protected from light, with fluorophore‑conjugated antibodies against CD4, CD8a, CD45RA, CD45RO, CCR7, CD25, PD‑1, CD137, CD69, and TIM‑3 (all from BioLegend; catalogue numbers listed in the Reagents and Resources table). All antibodies were diluted to 3% (v/v) in FACS buffer consisting of PBS supplemented with 0.5 mM EDTA (Invitrogen, Cat#10135423) and 2.5% (v/v) FBS (Gibco, Cat#17593595). Cells were washed and resuspended in FACS buffer.

For post‑co‑culture flow‑cytometry analysis, organoid–T cell co‑cultures and corresponding controls were dissociated into single‑cell suspensions by incubation with TrypLE Express (Gibco, Cat#12605028; 50 µL per well) combined with gentle mechanical disruption, followed by dilution with ice‑cold DPBS (Gibco, Cat#14190144). Samples were centrifuged at 600 × g for 5 minutes at 4 °C, and cell pellets were stained with Zombie Green Fixable Viability Dye (BioLegend, Cat#423112; 3 µL dye diluted in 1000 µL PBS, 50 µL applied per well) for 30 minutes at room temperature in the dark. After washing in FACS buffer and a subsequent centrifugation step (600 × g for 5 minutes at 4 °C), cells were stained (40 µL per well) for 30 minutes at 4 °C with fluorophore‑conjugated antibodies against CD4, CD8a, CD25, CD69, CD45, and CD326 (EpCAM) (all from BioLegend; catalogue numbers listed in the Reagents and Resources table). Each antibody was diluted to 3% (v/v) in FACS buffer.

All staining and wash steps were performed on ice and protected from light. Following antibody staining, cells were resuspended in 120 µL FACS buffer per well and analysed on a Novocyte Penteon flow cytometer (Agilent). At least 10,000 events were acquired for organoid single‑cell populations where applicable. Compensation was performed using UltraComp eBeads (Thermo Fisher Scientific, Cat#01‑2222‑42). Data were analysed using NovoExpress software version 1.6.2 (Agilent). Gating for singlets, live cells, organoids, and CD3+ T cell subsets was performed according to the strategy shown in Figure 2B. All flow‑cytometry experiments were performed with a minimum of three independent biological replicates, each with technical replicates, and results were compared between POC TCE–treated and matched control samples.

**Cytokines quantification**

At experimental endpoints, culture supernatants were collected, briefly centrifuged to remove cellular debris, and transferred to 96‑well V‑bottom plates. Samples were either processed immediately or stored at −80 °C until cytokine analysis. Concentrations of Granzyme B, Perforin, and Interferon‑γ (IFN‑γ) were quantified using the LEGENDplex Human CD8/NK Panel (13‑plex) with VbP V02 (BioLegend, Cat# 741187), according to the manufacturer’s instructions. Samples were acquired on a NovoCyte™ flow cytometer (Agilent), and FCS files were analysed using BioLegend LEGENDplex Data Analysis Software (https://legendplex.qognit.com). Data visualisation and statistical analyses were performed using GraphPad Prism version 10.4.1 (GraphPad Software).

**Antibody binding capacity (ABC) assay for EGFR and HER2 cell‑surface expression**

Cell‑surface expression of EGFR and HER2 on CRC organoids was quantified using an antibody binding capacity (ABC) assay employing the Quantibrite PE Phycoerythrin Fluorescence Quantitation Kit (BD Biosciences, Cat# 340495), according to the manufacturer’s instructions. Organoids were dissociated into single cells and seeded at 100,000 cells per well in 96‑well V‑bottom plates. Cell viability was assessed by staining with Zombie Green Fixable Viability Dye (BioLegend, Cat# 423112, 3 µL dye diluted in 1000 µL PBS, 50 µL applied per well) for 30 minutes at room temperature in the dark.

After washing, live cells were incubated for 1 hour at 4 °C in the dark with PE‑conjugated antibodies against human EGFR (BioLegend, Cat# 352904; 5 µL per 100 µL staining volume), human CD340 (ErbB2/HER2) (BioLegend, Cat# 324405; 5 µL per 100 µL staining volume), or a PE‑conjugated mouse IgG1 κ isotype control antibody (BioLegend, Cat# 400114; 5 µL per 100 µL staining volume).

Following antibody staining, cells were fixed with 4% (v/v) paraformaldehyde (PFA) in PBS for 20 minutes at room temperature in the dark, washed twice with FACS buffer as described above, and resuspended for acquisition. Quantibrite PE bead standards (BD Biosciences), consisting of four bead populations with defined, lot‑specific numbers of PE molecules per bead, were reconstituted in 500 µL PBS containing 0.5% (v/v) BSA and processed in parallel with samples according to the manufacturer’s instructions, without antibody staining.

Samples were acquired on a NovoExpress flow cytometer (Agilent), collecting >10,000 singlet live events per organoid sample. Quantibrite bead populations were identified using NovoExpress software version 1.6.2 (Agilent), and PE geometric mean fluorescence intensity (gMFI) values were used to generate standard curves by linear regression of gMFI versus the known number of PE molecules per bead, as specified for the corresponding bead lot. Absolute receptor expression was calculated by interpolating the PE gMFI of stained cells onto the Quantibrite standard curve, yielding antibody binding capacity values expressed as the number of antibody molecules bound per cell (molecules/cell). Specific antibody binding capacity values were obtained by subtracting the isotype‑matched ABC from the corresponding EGFR‑ or HER2‑specific ABC values. Measurements were performed on the indicated organoid models and compared between experimental conditions as specified.

**Organoid multiplex immunofluorescence (mIF) for EGFR and HER2 expression**

For multiplex immunofluorescence analysis, CRC organoids were dissociated into single cells and seeded at 4,000 cells per well in 96-well round-bottom plates pre-treated with Anti-Adherence Rinsing Solution (STEMCELL Technologies, Cat# 07010). Organoids were cultured in in-house CRC organoid medium (O.M.; lacking SB202190 and Y-27632) as described above, supplemented with 1% (v/v) UltiMatrix BME (Bio-Techne, Cat# BME001-10). Edge wells were filled with PBS to minimise evaporation.

Organoids were cultured in suspension for 5 days at 37 °C with 5% CO₂ prior to staining.

On the day of immunofluorescence analysis, culture medium was removed and organoids were gently washed with PBS. Fixation was performed by incubating samples with 4% paraformaldehyde (v/v) in PBS for 10 minutes at room temperature, followed by two washes with PBS under gentle agitation. For intracellular phospho‑EGFR detection, organoids were permeabilised with 0.3% (v/v) Triton X‑100 (Thermo Fisher Scientific, Cat# A16046.AP) in PBS for 10 minutes at room temperature and washed twice with PBS. Samples were blocked with 1% (v/v) bovine serum albumin (BSA) (Thermo Fisher Scientific, Cat# 15260037) in PBS for 1 hour at room temperature to minimise non‑specific antibody binding. Multiplex primary antibody staining was performed in blocking buffer using CoraLite Plus 488‑conjugated anti‑HER2/ErbB2 antibody (Proteintech, Cat# CL488‑60311; 1:100 dilution) and CoraLite Plus 647‑conjugated anti‑phospho‑EGFR (Tyr1197) antibody (Proteintech, Cat# CL647‑84906; 1:600 dilution). Antibody incubations were carried out for 1 hour at room temperature in the dark, followed by three PBS washes to remove unbound antibody. Nuclear counterstaining was performed using NucBlue Fixed Cell ReadyProbes DAPI reagent (Thermo Fisher Scientific, Cat# R37606) according to the manufacturer’s instructions.

Organoids were imaged using a BioTek Cytation5 Cell Imaging Multi‑Mode Reader (Agilent) equipped with brightfield, GFP, DAPI, and Cy5 filter cubes. All steps following antibody staining were protected from light. Stained plates could be stored at 4 °C in the dark for up to 24–48 hours prior to imaging. Multiplex immunofluorescence analyses of organoid models were carried out using Gen5 software version 3.16.10 (Agilent).

**Live‑cell motility and cytotoxicity imaging**

For live‑cell imaging assays assessing T cell motility, COLO250 organoids were cultured for 5 days in in-house CRC organoid medium (O.M.; lacking SB202190 and Y-27632) as described above, supplemented with 0%, 0.5%, 1%, 2.5%, 5%, or 100% (v/v) UltiMatrix BME (Bio‑Techne, Cat# BME001‑10) to evaluate the impact of matrix density on T cell movement. Organoids were subsequently co‑cultured for 72 hours with 7‑day pre‑stimulated CD3+ T cells from donor 054, which were labelled with CellTracker Deep Red Dye (Invitrogen, Cat# C34565) according to the manufacturer’s instructions. Co‑cultures were maintained in a BioTek BioSpa 8 Automated Incubator (Agilent), and time‑lapse imaging was performed every 12 hours using a BioTek Cytation 5 Cell Imaging Multi‑Mode Reader (Agilent) equipped with brightfield and Cy5 filter cubes at 10× magnification. For cytotoxicity imaging, organoids were dissociated into single cells, labelled with 5 µM Incucyte Caspase‑3/7 Green Dye for Apoptosis (Sartorius, Cat# 4440), and cultured for 5 days in organoid medium supplemented with 1% (v/v) UltiMatrix BME (Bio‑Techne, Cat# BME001‑10). Organoids were then co‑cultured for 72 hours with 7‑day stimulated CD3+ T cells from donor 054, labelled with CellTracker Deep Red and treated with T cell engager molecules at the indicated concentrations. Time‑lapse imaging was performed as described above using brightfield, GFP, DAPI, and Cy5 filter cubes. No medium exchange was performed during the imaging period. Image acquisition and quantitative analysis were carried out using Gen5 software version 3.16.10 (Agilent).

**Genomic DNA and RNA extraction, library preparation, and sequencing**

Genomic DNA and total RNA were isolated from differentiated CRC organoid pellets using the Monarch Genomic DNA Purification Kit (NEB, Cat#T3010) and the Monarch Spin RNA Isolation Kit (Mini) (NEB, Cat#T2110), respectively, following the manufacturer’s instructions. Organoid pellets were processed immediately or stored at −80 °C prior to nucleic acid extraction. DNA and RNA concentration, fragment size distribution, and integrity were assessed using the TapeStation 4200 system (Agilent).

For whole‑genome sequencing, genomic DNA was sheared using a Covaris Sonicator (Covaris, Cat#E220) and g‑TUBEs (Covaris, Cat#520079) according to the manufacturer’s recommendations, followed by library preparation using the Ligation Sequencing Kit V14 (Oxford Nanopore Technologies, Cat#SQK‑LSK114).

For transcriptome analysis, total RNA was used directly for library preparation using the Direct RNA Sequencing Kit (Oxford Nanopore Technologies, Cat#SQK‑RNA002). Prepared libraries were loaded onto PromethION Flow Cells Packs (DNA: ONT Cat#FLO‑PRO114M; RNA: ONT Cat#FLO‑PRO004RA) and sequenced according to standard Oxford Nanopore Technologies protocols on a PromethION 24 A100 platform (ONT).

**SUPPLEMENTARY FIGURES**

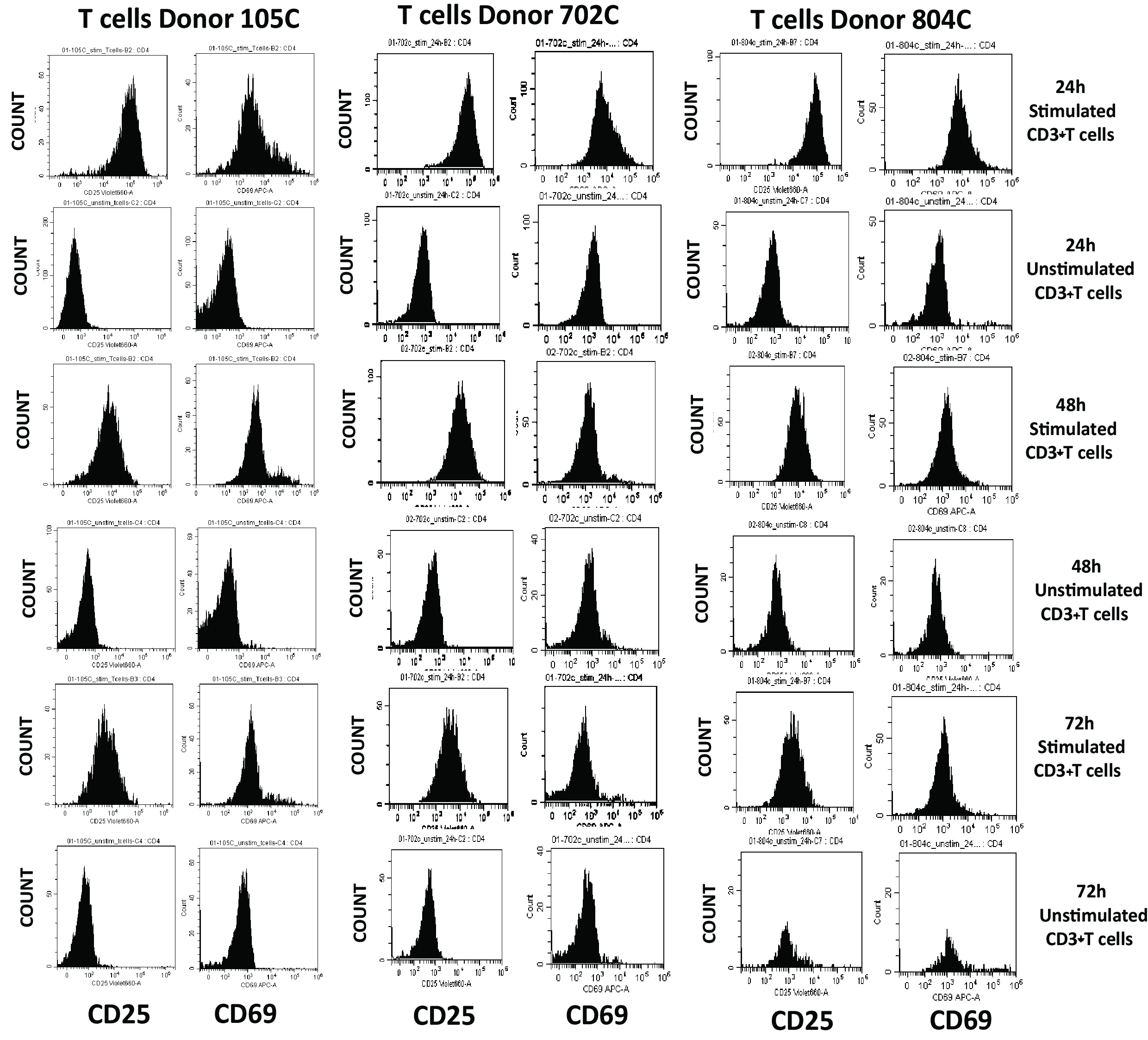

**Figure S1.** **CD3+ T cell activation following isolation (Related to Figure 1).** Single-parameter flow cytometry histograms showing CD25 and CD69 expression in CD3+T cells from leukopak donors 105C, 702C, and 804C at 24, 48, and 72 h post-isolation. Fluorescence intensity distributions are shown after sequential gating on singlets, live cells, and CD3+T cells, with unstained and isotype controls used to define background signal. These data were used to assess activation kinetics and donor-to-donor variability prior to co-culture experiments. Representative data are shown from three independent biological experiments, each performed with technical replicates.

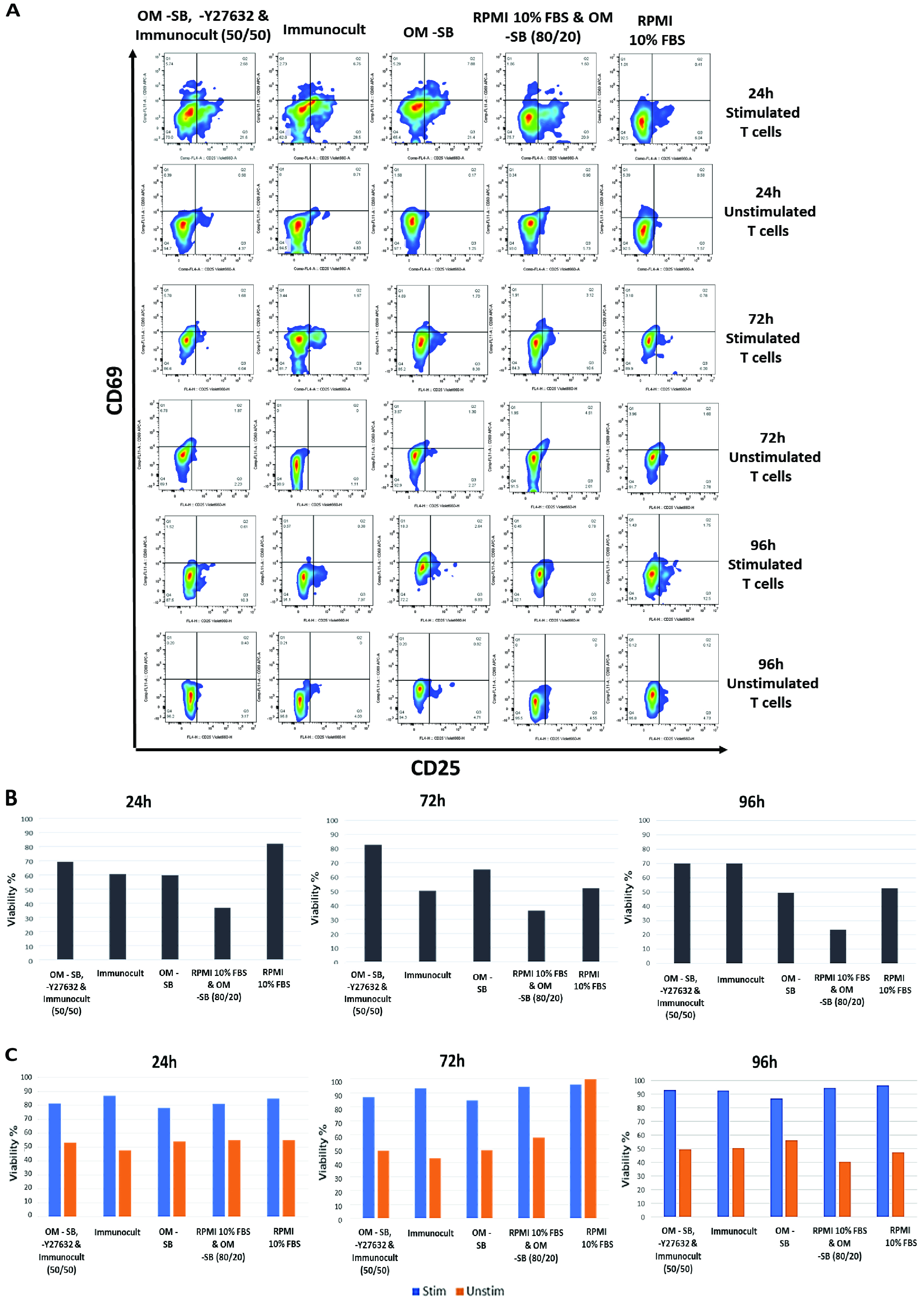

**Figure S2. Organoid–CD3+ T cell co**‑**culture and controls conditions across media formulations (Related to Figure 1).** Organoid–CD3+ T cell co‑cultures and matched control conditions were assessed under different media formulations at 24, 72, and 96 h. **(A)** Activation of stimulated and unstimulated CD3+ T cells in co‑culture with organoids under the indicated media conditions. **(B)** Viability of organoid‑only control cultures over time. **(C)** Viability of stimulated and unstimulated CD3+ T cell‑only control cultures over time. Viability was measured via flow cytometry viability staining. Values were normalized to the respective organoid model as mono-culture at the indicated time point. Data represent mean ± SD from at least three independent biological experiments, each performed with technical replicates. O.M., organoid medium; SB, SB202190; Stim, stimulated CD3+T cells; Unstim, unstimulated CD3+T cells.

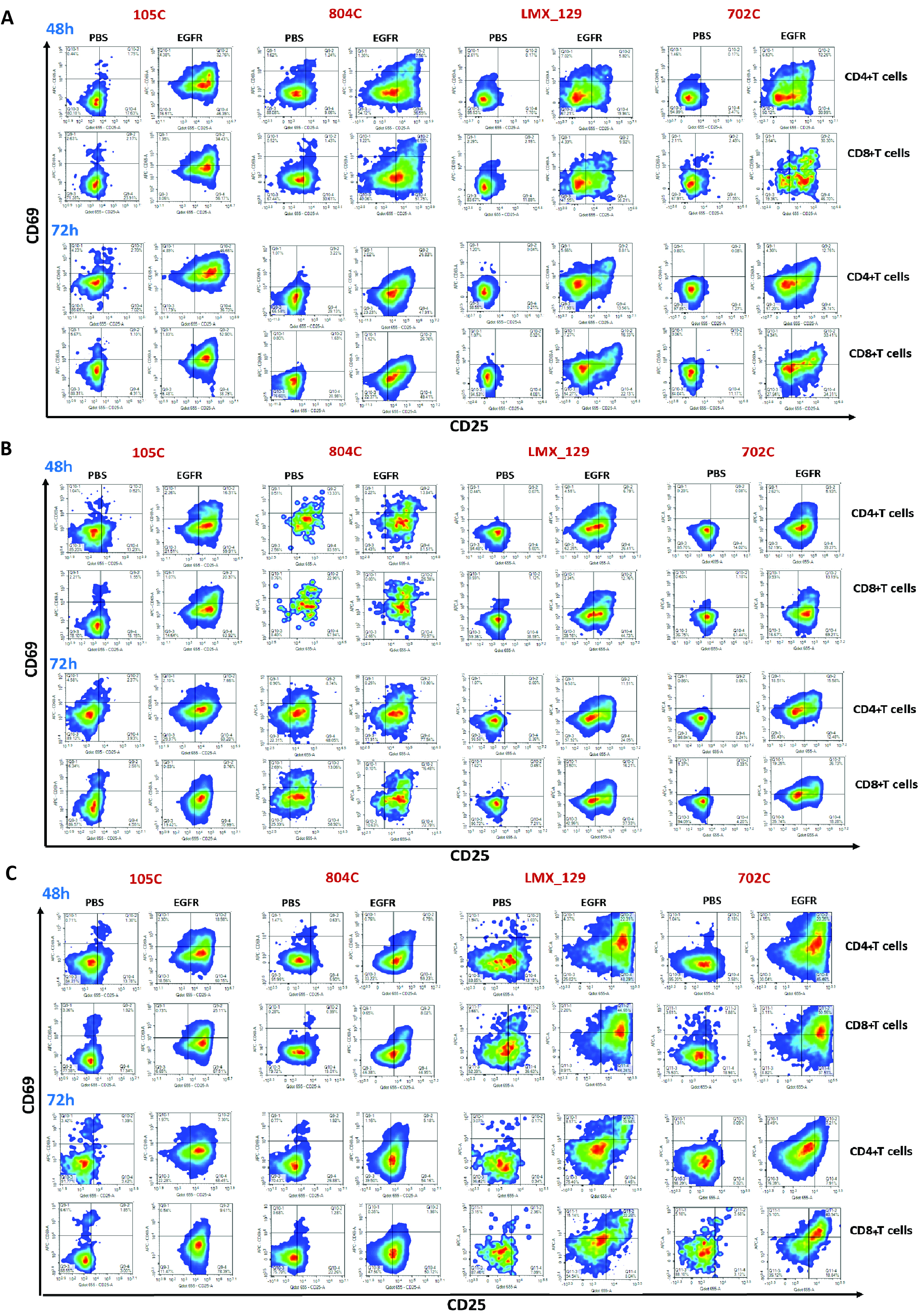

**Figure S3. EGFR**‑**TCE–induced CD4 and CD8+T cells activation in CRC organoid co**‑**cultures (Related to Figure 3).** CRC organoids (COLO250, COLO298, and S315571) were co‑cultured with stimulated CD3+T cells from healthy donors (105C, 804C, 702C, and LMX_129) and treated with EGFR‑TCE (100 nM) or PBS for 48 h and 72 h. (**A**) Activation of stimulated CD4 and CD8+ T cell in co‑culture with COLO250 organoids. (**B**) Activation of stimulated CD4 and CD8+ T cell in co‑culture with COLO298 organoids. (**C**) Activation of stimulated CD4 and CD8+ T cell in co‑culture with S315571 organoids. T cell activation was quantified by flow cytometry based on CD25 and CD69 expression following sequential gating as described in **Fig. 2B**. Data represent mean ± SD from at least three independent biological experiments, each performed with technical replicates.

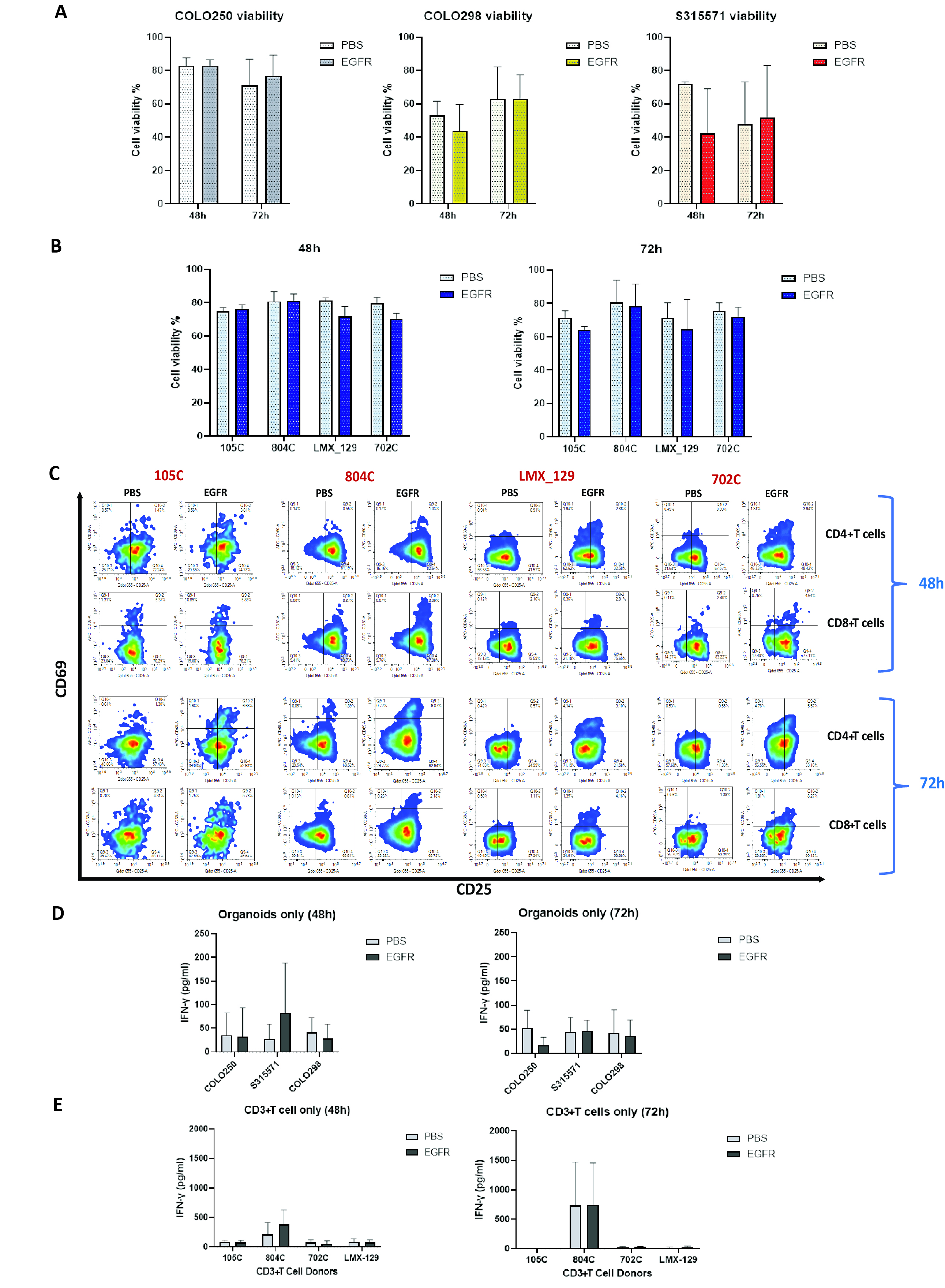

**Figure S4. CRC organoid and T-cell-only control cultures following treatment with EGFR-TCE (100 nM) or PBS for 48 or 72 h (related to Figure 3).** CRC organoids (COLO250, COLO298, and S315571) and stimulated CD3+ T cells from healthy donors (105C, 804C, 702C, and LMX_129) were cultured separately and treated with EGFR-TCE (100 nM) or PBS. **(A)** Viability of CRC organoid-only cultures. **(B)** Viability of stimulated CD3+ T-cell-only cultures. **(C)** Activation of CD4+ and CD8+ T cells from donors 105C, 804C, 702C, and LMX_129. **(D)** IFN-γ concentrations in supernatants from organoid-only and T cell-only control cultures. Data represent mean ± SD from at least three independent biological experiments, each performed with technical replicates.

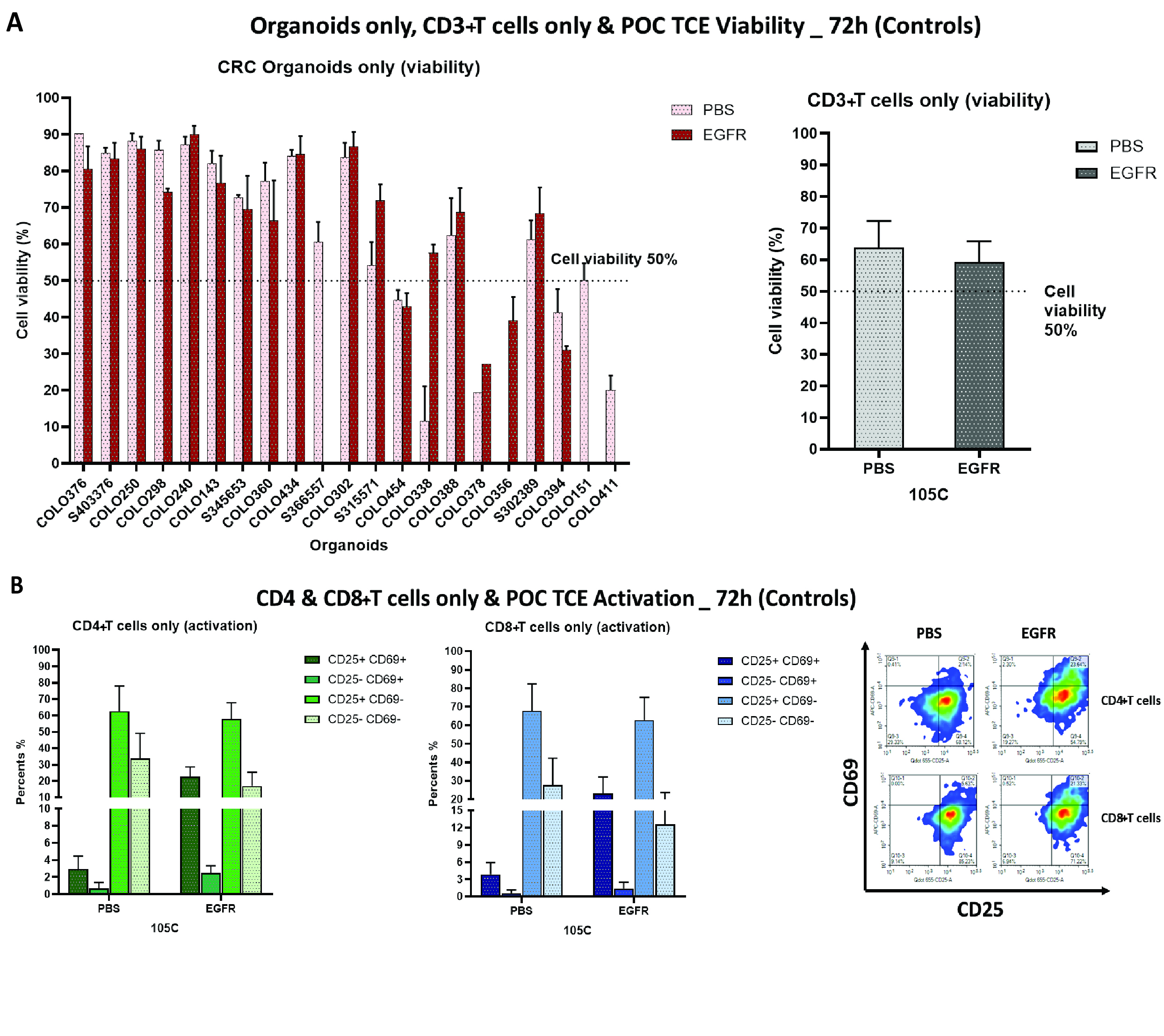

**Figure S5.** **CRC organoids and stimulated CD3+ T cells (105C donor) only control following treatment with EGFR-TCE (100 nM) or PBS for 72 h (related to Figure 4).** CRC organoids and stimulated CD3+ T cells from donor 105C were cultured separately and treated with EGFR-TCE (100 nM) or PBS. **(A)** Viability of 21 CRC organoid-only cultures. **(B)** Viability of stimulated CD3+ T cells-only cultures. **(C)** Activation of stimulated CD4+ and CD8+ T cells cultured in the absence of organoids. Data represent mean ± SD from at least three independent biological experiments, each performed with technical replicates.

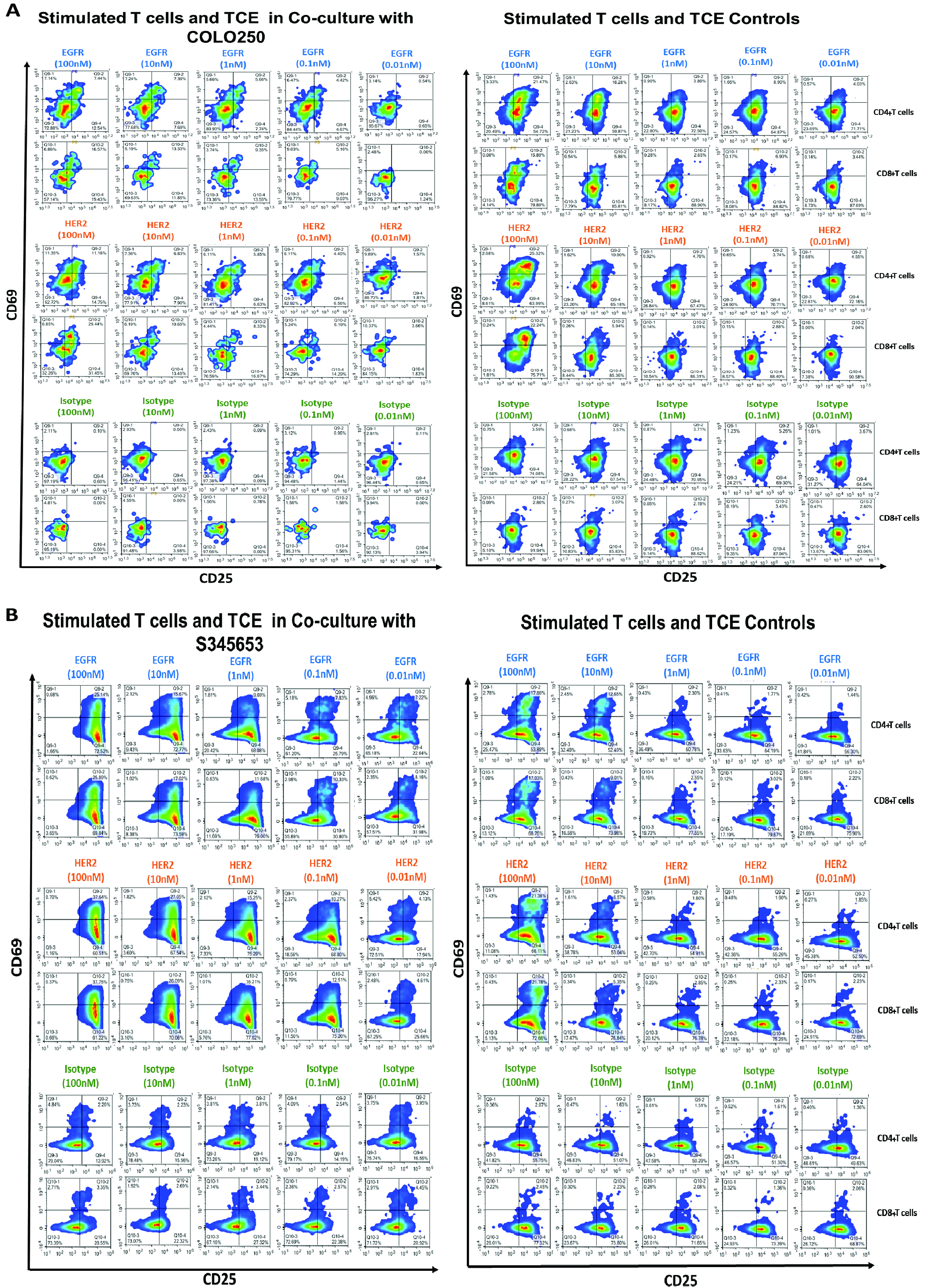

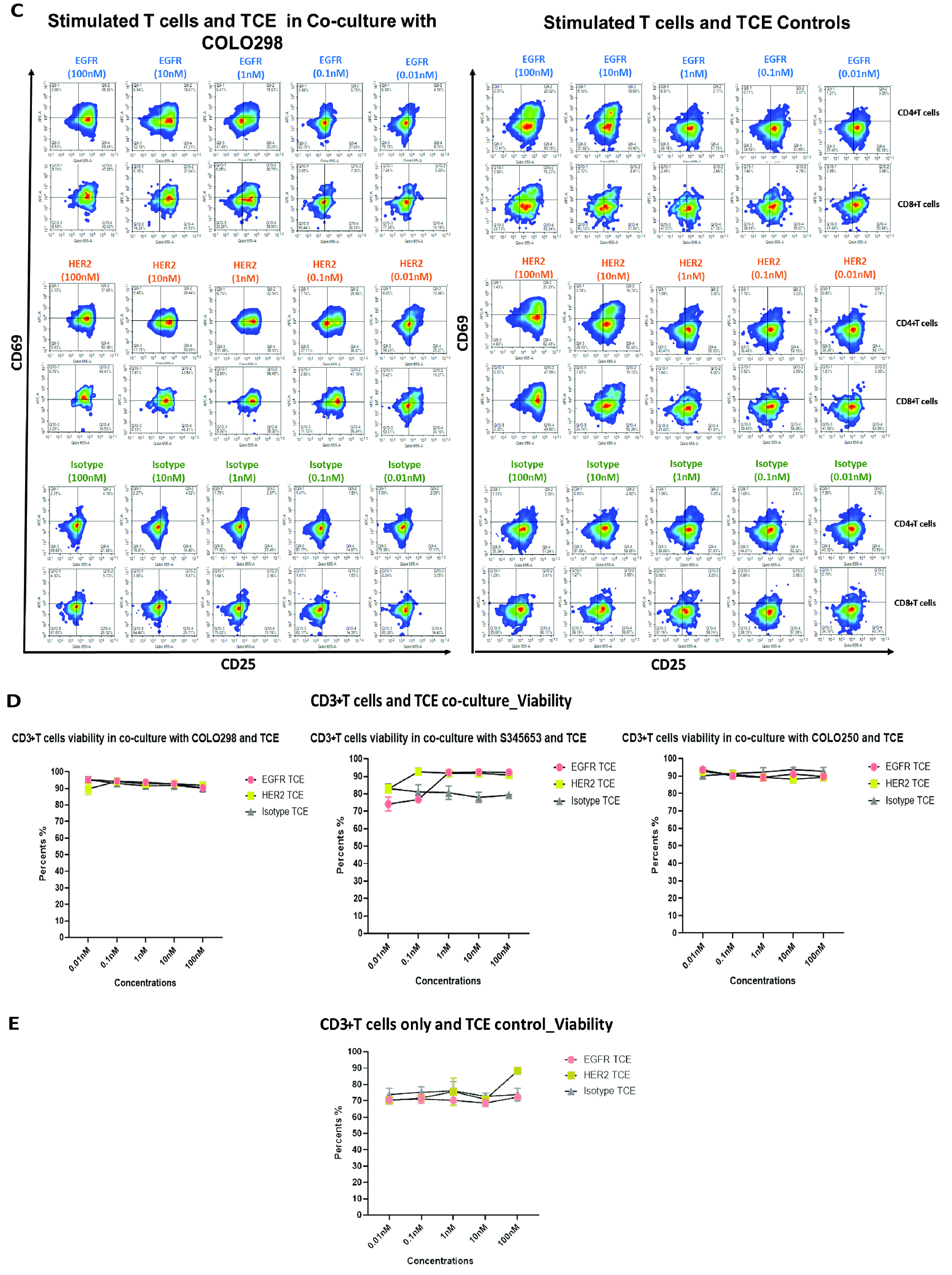

**Figure S6. POC-TCE dose-response effects in CRC organoid-T-cell co-cultures (Related to Figure 6).** CRC organoids (COLO250, S345653, and COLO298) were co-cultured with stimulated CD3+ T cells from donor 105C and treated with increasing concentrations of POC-TCE molecules for 72 h. **(A)** Activation of CD4+ and CD8+ T cells following co-culture with COLO250 organoids and corresponding T cell-only controls.**(B)** Activation of CD4+ and CD8+ T cells following co-culture with S345653 organoids and corresponding T-cell-only controls. **(C)** Activation of CD4+ and CD8+ T cells following co-culture with COLO298 organoids and corresponding T cell-only controls. T cell activation was assessed by flow cytometry based on CD25 and CD69 expression. **(D)** Viability of organoid-only control cultures treated with POC-TCE molecules. **(E)** Viability of stimulated CD3+ T cell-only control cultures treated with POC-TCE molecules. Viability was measured via flow cytometry viability staining. Values were normalized to the respective organoid model as mono-culture at the indicated time point. Data represent mean ± SD from at least three independent biological experiments, each performed with technical replicates.

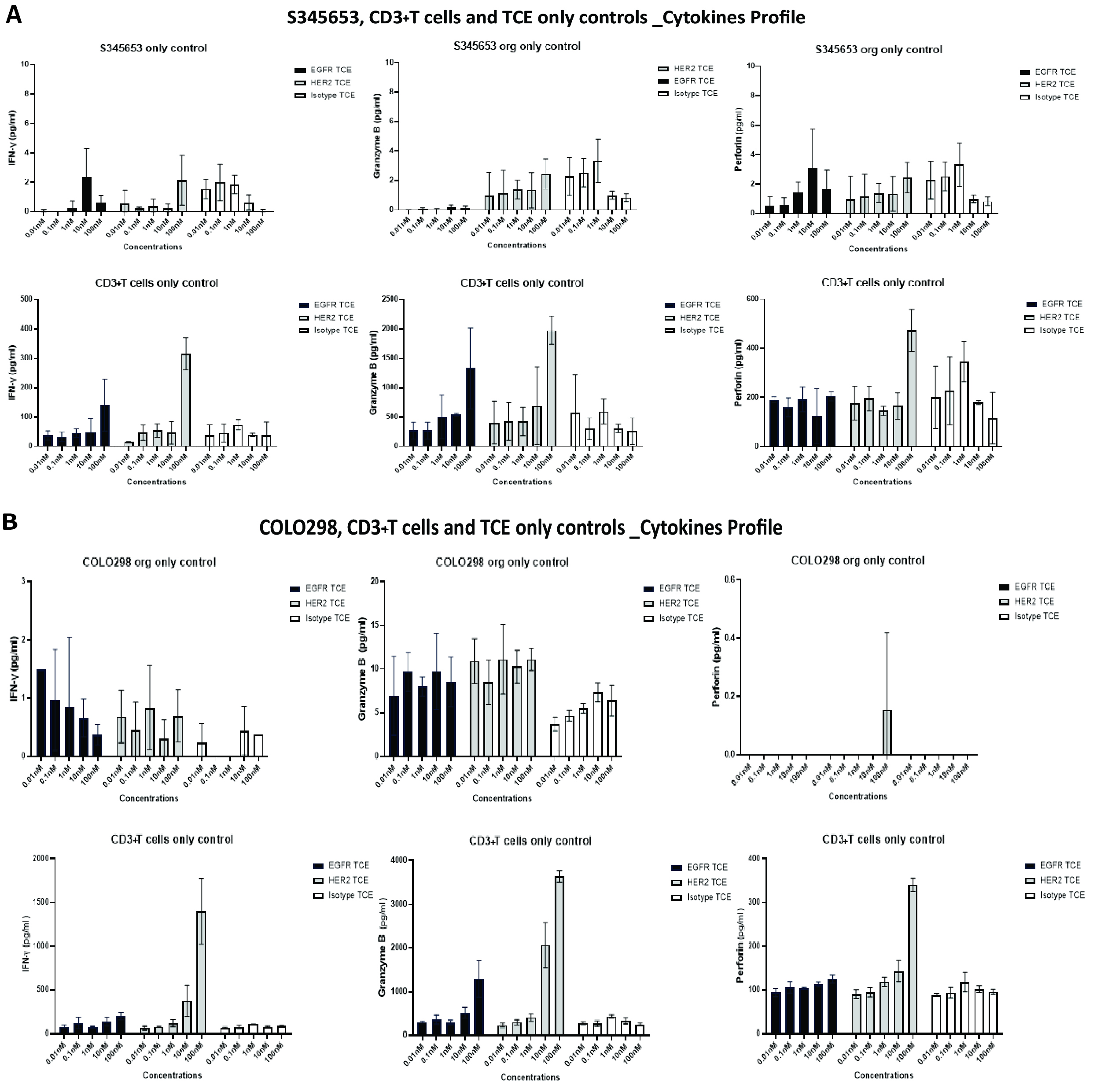

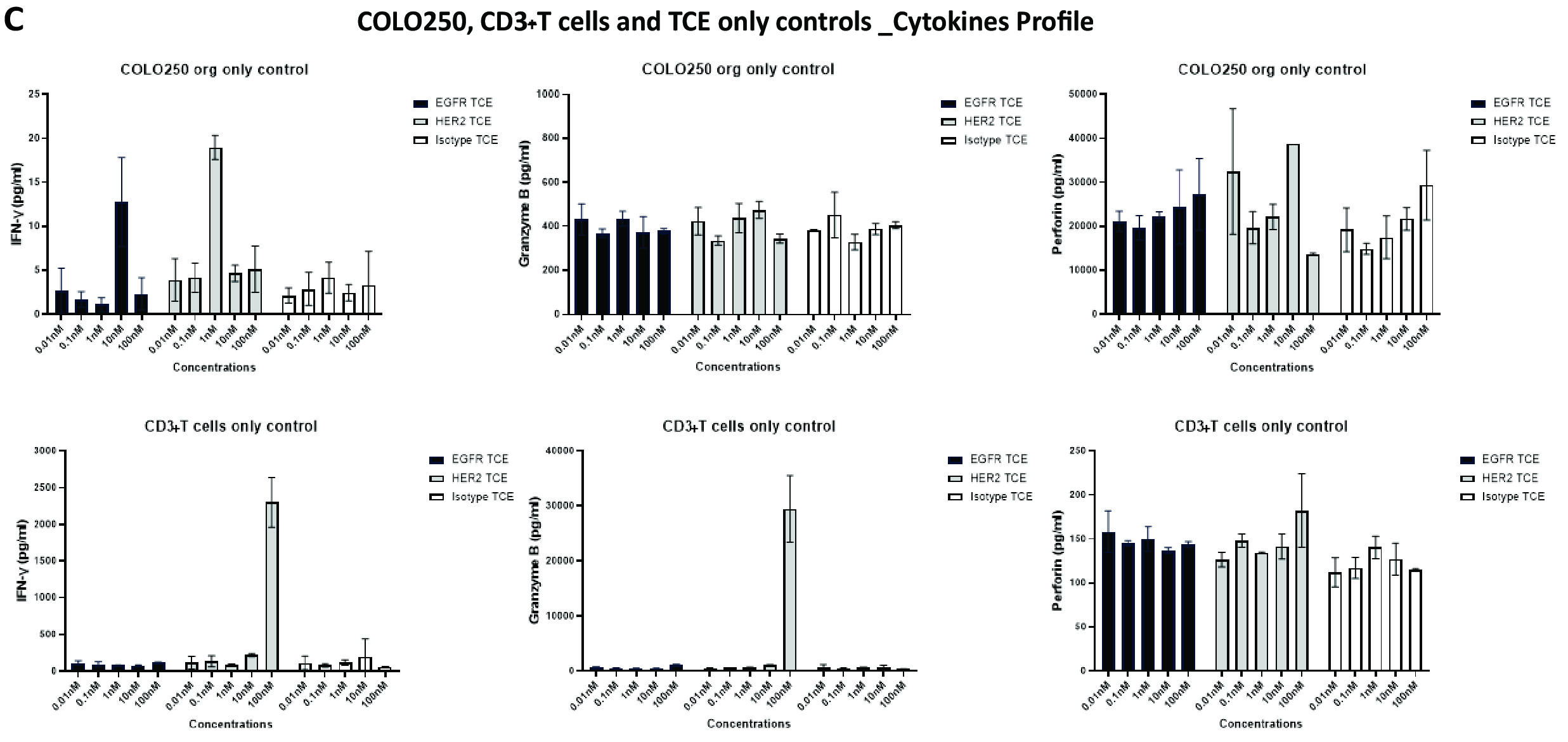

**Figure S7. Cytokine secretion following treatment with POC-TCE molecules in CRC organoid and T cell-only cultures (related to Figure 6).** Cytokine concentrations were quantified after 72 h of treatment with POC-TCE molecules in CRC organoid-only cultures (top panels) and matched activated CD3+ T cell-only control cultures (donor 105C, bottom panels). **(A)** IFN-γ, granzyme B, and perforin concentrations in S345653 organoid-only cultures and matched T cell-only controls. **(B)** IFN-γ, granzyme B, and perforin concentrations in COLO298 organoid-only cultures and matched T cell-only controls. **(C)** IFN-γ, granzyme B, and perforin concentrations in COLO250 organoid-only cultures and matched T cell-only controls. Cytokine concentrations are displayed using scales optimised for each organoid model to facilitate comparison of treatment-dependent effects within each organoid or matched T cell-only control condition rather than direct comparison of absolute cytokine levels between models. Data represent mean ± SD from three independent biological experiments, each performed with technical replicates.
